# GSK3 and Lamellipodin balance lamellipodial protrusions and focal adhesion maturation in mouse neural crest migration

**DOI:** 10.1101/2022.12.23.521694

**Authors:** Lisa Dobson, William B. Barrell, Zahra Seraj, Steven Lynham, Sheng-Yuan Wu, Matthias Krause, Karen J. Liu

## Abstract

Neural crest cells are multipotent cells that delaminate from the neuroepithelium, migrating to distant destinations throughout the embryo. Aberrant migration has severe consequences, such as congenital disorders. While animal models have improved our understanding of neural crest anomalies, the *in vivo* contributions of actin-based protrusions are still poorly understood. Here, we demonstrate that murine neural crest cells use lamellipodia and filopodia *in vivo*. Using neural crest-specific knockouts or inhibitors, we show that the serine-threonine kinase, Glycogen Synthase Kinase-3 (GSK3), and the cytoskeletal regulator, Lamellipodin (Lpd), are required for lamellipodia formation whilst preventing focal adhesion maturation. We consequently identified Lpd as a novel substrate of GSK3 and found that phosphorylation of Lpd favours Lpd interactions with the Scar/WAVE complex (lamellipodia formation) at the expense of Ena/VASP protein interactions (adhesion maturation and filopodia formation). All together, we provide an improved understanding of cytoskeletal regulation in mammalian neural crest migration, which has general implications for neural crest anomalies and cancer.

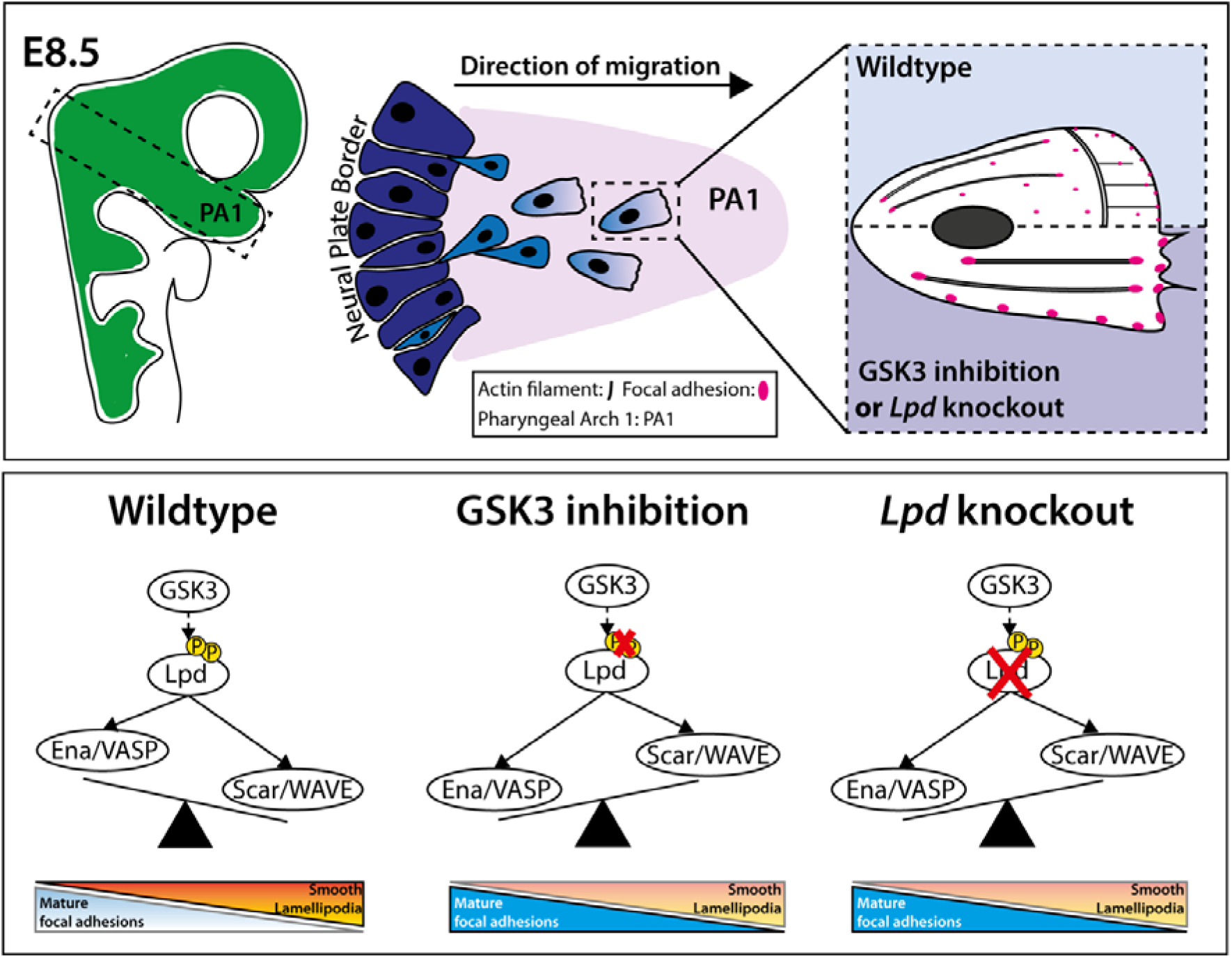

## INTRODUCTION

Neural crest cells are highly migratory multipotent cells that give rise to diverse tissues, such as the craniofacial skeleton and the peripheral nervous system^1^. Following their specification and induction, neural crest cells delaminate from the neural plate border, undergo an epithelial-to-mesenchymal transition (EMT), and migrate long distances to populate distant organs (for review, see: ^1–3^). In *Xenopus,* chicken and mouse, cranial neural crest cells migrate within defined streams towards the frontonasal process (FNP) and pharyngeal arches (PA) 1 and 2 ^2,4–7^.

Mesenchymal migration requires the formation of actin-based protrusions at the cell leading edge which are constantly interacting with the surrounding extracellular matrix (ECM) through integrin-based adhesions^8–12^. Because the *in vivo* migration of neural crest cells is highly species-specific^2,13,14^, it is unclear whether the use of specific actin-based protrusions, such as sheet-like protrusions (lamellipodia) or finger-like protrusions (filopodia), is conserved. We and others have proposed that mouse cranial neural crest cells use lamellipodial protrusions for efficient locomotion *in vitro*^13^. However, due to the difficulty in live imaging mouse neural crest cells, very little is known about the cytoskeletal requirements for *in vivo* migration.

We previously showed that the serine-threonine kinase Glycogen Synthase Kinase-3 (GSK3) is required during mouse neural crest development^13^. Conditional loss of both isoforms (GSK3α and GSK3β) led to a loss of expression of the neural crest-specific transcription factor, *Sox10,* in the facial prominences, with an associated failure of neural crest migration^13^. Pharmacological inhibition of GSK3 caused a collapse of lamellipodia^13^, raising the intriguing possibility that GSK3 may act via Lamellipodin (Lpd, HUGO name: RAPH1). Lpd is an actin regulator that is an effector of the small GTPase Rac1, the key regulator of lamellipodia and mesenchymal cell migration^10,15,16^. Indeed, loss of GSK3 activity in cultured mouse neural crest cells led to mislocalisation of Rac1 and Lpd^13^. Here, we report that the *in vivo* deletion of *Lpd* in neural crest cells mimics the cellular effects of GSK3 inhibition, with an increase in filopodial protrusions.

Lpd promotes cell migration via interactions with both the Scar/WAVE complex and with Ena/VASP proteins^16,17^. The Scar/WAVE complex is essential for lamellipodia formation and is composed of five proteins (Sra1/Pir121, Nap1, Scar/WAVE1-3, Abi1-3, and HSPC300), which activate the Arp2/3 complex to induce branched actin filament nucleation^10,18,19^. In contrast, Ena/VASP proteins (VASP, Mena, EVL) increase actin filament length at the cell leading edge by temporarily preventing capping of actin filaments and recruiting polymerisation-competent G-actin bound to profilin^20–27^. In addition, Ena/VASP proteins are required for integrin-based adhesion maturation^28^.

Here, we show that mouse neural crest cells use lamellipodia and filopodia for migration. Loss of *Lpd* caused a switch from lamellipodial to filopodial protrusion *in vitro* and *in vivo.* The cell behaviours are reminiscent of the published Arp2/3 knockouts whereby Wu and colleagues (2012)^29^ examined mouse embryonic fibroblasts (MEFs). This suggests that neural crest cell migration depends on Lpd functioning via Scar/WAVE-Arp2/3 complexes. In addition, we found that GSK3 acts via Lpd to inhibit focal adhesion maturation underneath the lamellipodium. Increased GSK3 kinase activity promoted Lpd interaction with the Scar/WAVE complex while reducing interaction with the Ena/VASP proteins, VASP and Mena. We then used mass spectrometry analysis to identify multiple Ser/Thr GSK3-dependent phosphorylation sites in Lpd, including sites localised within the C-terminus that overlap with known Scar/WAVE complex and Ena/VASP protein binding sites^15,16^. Thus, GSK3 and Lpd cooperate to coordinate lamellipodial protrusions with adhesion maturation to support actin-based migration of mouse cranial neural crest cells.

## RESULTS

### Murine neural crest cells use lamellipodia and filopodia in vivo

Mouse neural crest cells are known to undergo defined, directed migration. However, a detailed analysis of their *in vivo* membrane protrusions has not previously been performed. To do this, we used mouse lines carrying either Cre-dependent *LifeAct-EGFP* or *R26R^mtmg^*. *LifeAct-EGFP* encodes a reporter of filamentous actin dynamics, while *R26R^mtmg^* animals express membrane tagged fluorescent Tomato (mT), which is switched to membrane green fluorescent protein (mGFP) upon breeding with a Cre-transgene^30,31^. Here, we labelled neural crest cells by inter-crossing either reporter line to the neural crest-specific *Wnt1::cre* line^32^.

In wildtype embryos, mGFP-labelled cranial neural crest cells (*Wnt1::*cre; *Rosa26R^mtmg^)* can be visualised by the 4 somite stage (4ss) at approximately embryonic day 8.5 (E8.5) (Figure 1A). By the 7-somite stage (Figure 1B), neural crest cells delaminate from the neuroepithelium and migrate toward the pharyngeal arches (Figure 1A-B). At this stage, we can observe cells destined for pharyngeal arch 1 (PA1), which will give rise to the jaws (Fig 1C, yellow arrowhead). Similarly, migration of the 2^nd^ pharyngeal arch (PA2) stream is also under way (Figure 1C, white arrowhead), whilst the vagal/cardiac neural crest cells can be seen emerging (Figure 1C, asterisk), as schematised in (Figure 1D).

**Figure 1.**
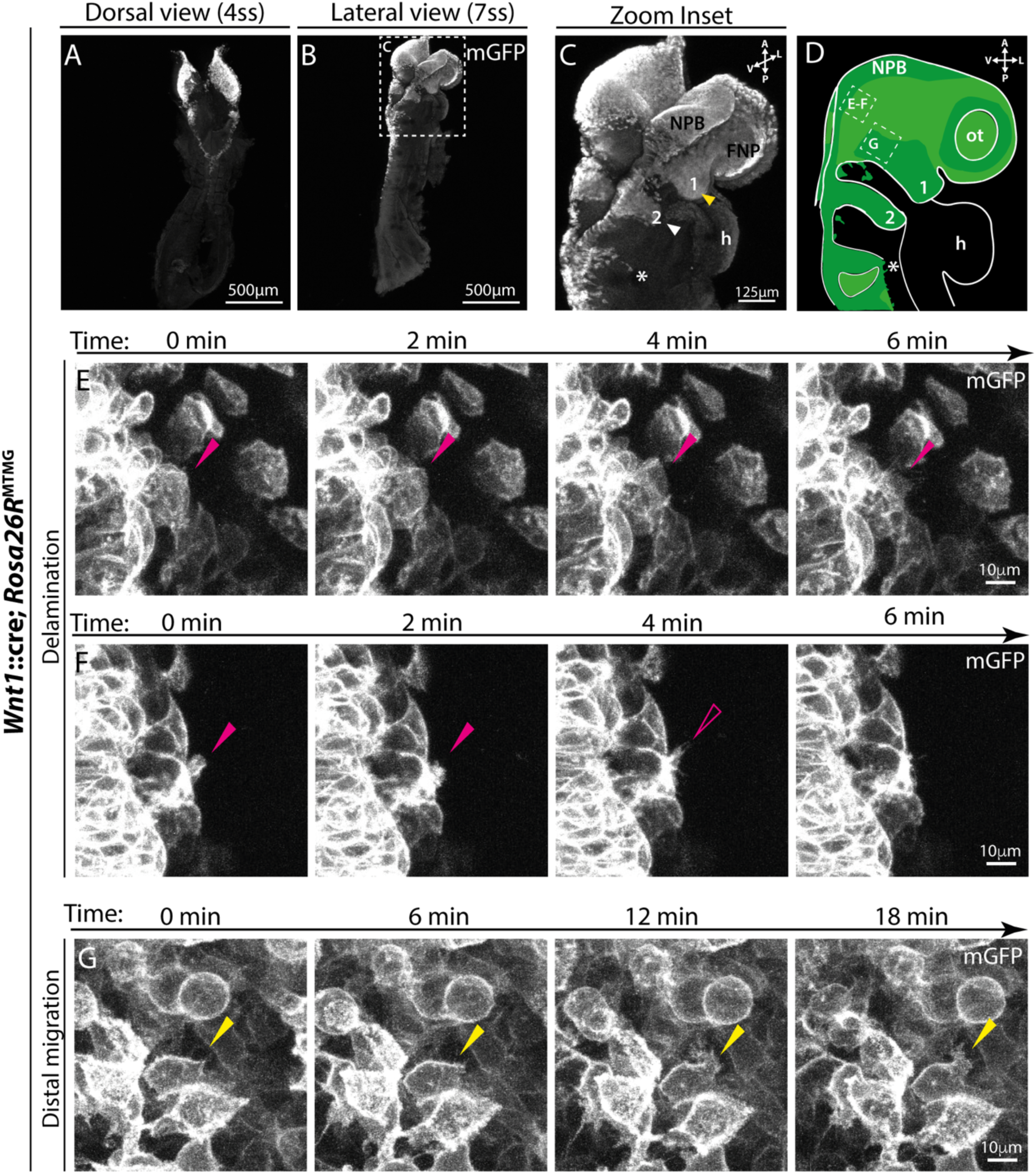
Mouse cranial neural crest cells display lamellipodia and filopodia *in vivo*. **(A-B)** Maximum projection images of E8.5-8.75 mouse embryos, with the neural crest lineage-labelled with mGFP (*Wnt1::*cre; *Rosa26R^mtmg^)*, shown as **(A)** dorsal view (4 somite stage, ss) and **(B)** lateral view (7ss). Scale bar: 500 µm. **(C)** Zoom inset of **(B)** showing the cranial neural crest streams of an E8.75 embryo (7ss). Cranial neural crest cells leave the neural plate border (NPB) and migrate to populate the frontal nasal process (FNP), pharyngeal arch 1 (PA1, yellow arrowhead) and pharyngeal arch 2 (PA2, white arrowhead). Vagal and cardiac neural crest streams also start to emerge (*). Scale bar: 125 µm. **(D)** Schematic representation of a laterally-oriented E8.75 embryo (7-9ss), with white dashed boxes showing the imaging regions of interest (ROIs) in **(E-G)**. Green: lineage-labelled neural crest streams. h: heart, ot: otic vesicle. **(E-G)** Time-lapse stills from live imaging of laterally-oriented wild-type (*Wnt1::*cre; *Rosa26R^mtmg^)* E8.75 embryos, with neural crest cells labelled with mGFP. **(E-F)** Time-lapse stills from two example movies of delaminating cranial neural crest (solid magenta arrowhead: lamellipodia and open magenta arrowhead: filopodia). 10 minute movies (1 frame/ 20 seconds), z-depth 24.5 µm, 0.5 µm per slice. See **Supplementary Movies 1-2**. **(G)** Filopodial protrusions on distally migrating cranial neural crest cells (yellow arrowhead). 30 minute movie (1 frame/ 45 seconds), z-depth 40 µm, 0.5 µm per slice, see **Supplementary Movie 3**. Movies are representative of 3 embryos, all 6-10 somite stage (n=3). Scale bar: 10 µm.

To visualise the membrane protrusions of individual cranial neural crest cells destined for PA1, we used live confocal microscopy to image these cells *in vivo* (Figure 1E-G). We observed that migratory neural crest cells have cellular protrusions interacting with other neural crest cells as well as with surrounding non-neural crest mesenchyme and the extracellular environment (Supplementary Movies 1-3). Whilst most protrusions observed were filopodial, lamellipodia were seen emanating from delaminating cranial neural crest cells (Figure 1E (magenta arrowhead), Supplementary Movie 1). In some instances, lamellipodia resolved into filopodia following initial outward protrusion (Figure 1F (unfilled magenta arrowhead), Supplementary Movie 2). In contrast, in more distal domains, broad protrusions were infrequently observed (Figure 1G). Instead, multiple filopodia were seen extending from the edge of broad protrusive structures (Figure 1G (yellow arrowhead) and Supplementary Movie 3).

### Glycogen Synthase Kinase-3 (GSK3) is required for lamellipodia formation and neural crest migration

We previously reported a collapse of neural crest lamellipodia upon pharmacological inhibition or genetic knockout of both GSK3 isoforms (*GSK3*α/*GSK3*β) in mouse, which led to inefficient cell migration^13^. How GSK3 regulates lamellipodia in mouse neural crest cells, and whether this affects the cell speed and persistence at the single cell level remains unknown. To address this, we used neural crest explant cultures, where we dissect out the neural plate border at E8.5, during the early stages of neural crest induction (Figure 2A-B). This allows systematic assessment of mouse neural crest cells as they delaminate and migrate away from the neural plate border (Figure 2C)^33^. Neural crest cells can then be treated with pharmacological inhibitors at specific time points, for example prior to delamination or during cell migration. Here, we cultured our wildtype explants for 24 hours and treated neural crest cells (analogous to E9+ migratory cells) with two different GSK3 pharmacological inhibitors (either BIO or CHIR99021), or DMSO, the vehicle control. Cells were pre-treated for 1 hour, then continuously imaged live for 18 hours (Figure 2D-F, Supplementary Movie 4). Cells were tracked throughout the course of time-lapse imaging to generate trajectory plots (Figure 2D’-F’).

**Figure 2.**
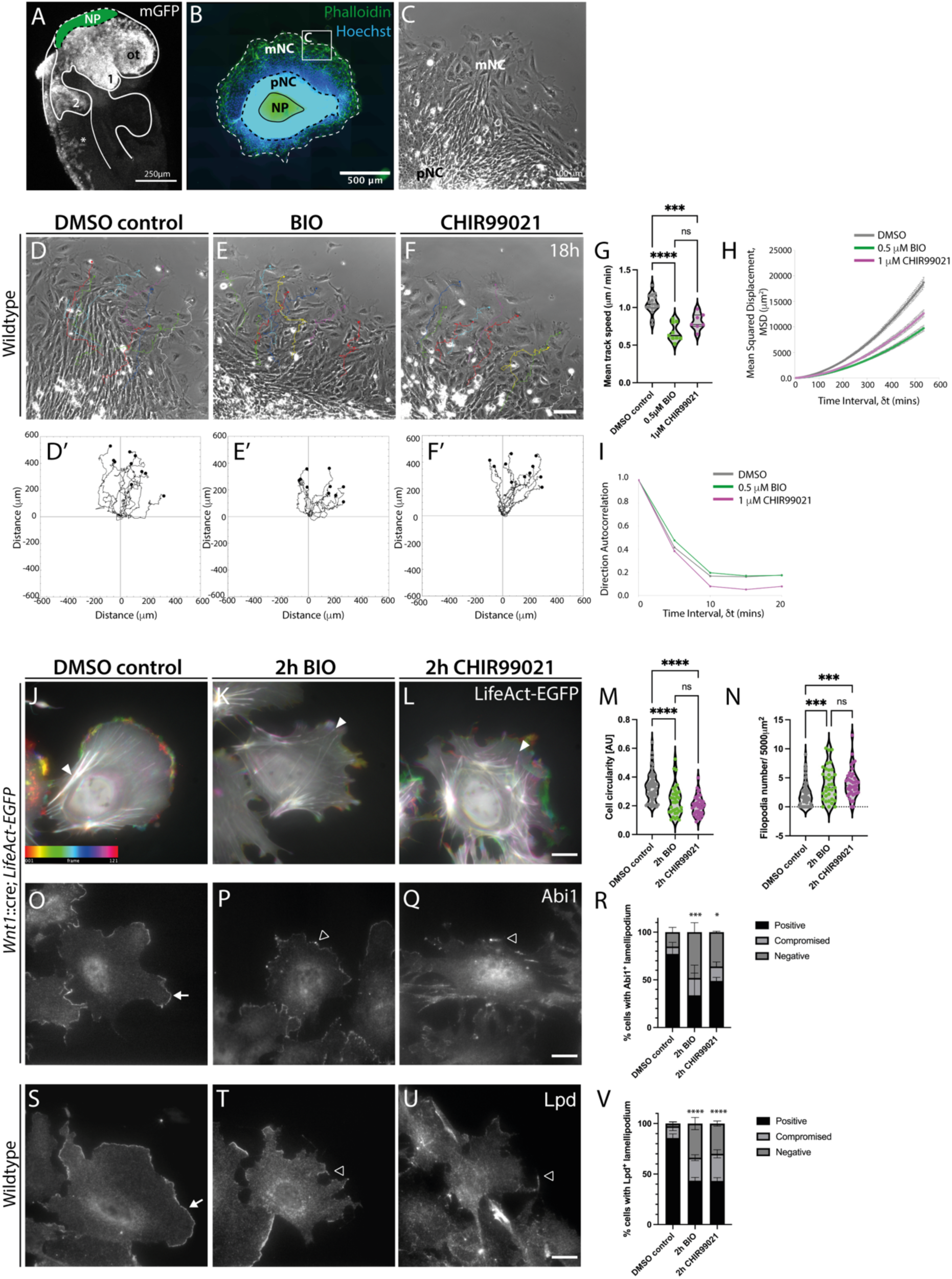
Inhibition of GSK3 prevents lamellipodia formation and reduces the migration efficiency of cranial neural crest cells *ex vivo*. **(A-C)** Neural crest explant cultures. **(A)** Schematic representation of a laterally-oriented *Wnt1::cre; Rosa26R“’mg* E8.5 embryo (5-8 ss), with neural crest cells lineage-labelled with mGFP and the dissected neural plate border (NP) pseudo-coloured green. 1: pharyngeal arch 1 (PA1), 2: PA2, ot: otic vesicle. **(B)** Neural crest explant culture 24 hours after dissection, with the actin filaments stained with phalloidin (green) and nuclei labelled with Hoechst (blue). The dissected NP is surrounded by pre-migratory neural crest cells (pNC), and an outer migratory neural crest population (mNC). Box represents imaging area used, as in **(C)**. Scale bar: 500 µm. **(C)** Phase-contrast image of the neural crest explant edge, used for *ex vivo* cell phenotyping. Scale bar: 100 µm. **(D-F)** Representative stills from 10x magnification phase-contrast time-lapse imaging of neural crest explants, cultured from E8.5 wildtype embryos treated with **(D)** DMSO vehicle control, or the GSK3 inhibitors, **(E)** 0.5 µM BIO or **(F)** 1 µM CHIR99021 for 18 hours (1 frame/ 5 min) (see Supplementary Movie 4). Scale bar 100 µm. **(D’-F’)** Trajectory plots of 10 migratory neural crest cells tracked through time-lapse imaging for **(D-F)**. **(G-I)** Quantification of **(G)** Mean Track Speed (MTS), **(H)** Mean Squared Displacement (MSD) and (I) direction autocorrelation of migratory cranial neural crest cells. For MTS calculations, each dot represents the mean speed of 10 neural crest cells imaged from the same explant: DMSO n=17 explants, 170 cells tracked; BIO n=8 explants, 80 cells tracked; CHIR99021 n=7 explants, 70 cells tracked, taken over 4 independent experiments. **** p < 0.0001, *** p < 0.001, ns non-significant, one-way ANOVA, Tukey’s multiple comparisons test. For direction autocorrelation measures, ∂t = 1, TR = 4 (20 min); see Materials & Methods for details. (K-M) Colour-coded time projection stills of 60x magnification time-lapse imaging (10 min movies, 1 frame/ 5 seconds) of migratory neural crest cells, cultured from E8.5 *Wnt1*::cre; *LifeAct-EGFP* embryos treated with (J) DMSO control, (K) 0.5 µM BIO or (L) 1 µM CHIR99021, 2h prior to imaging (see Supplementary Movie 5). Scale bar 20 µm. (M-N) Quantification of neural crest (M) cell circularity and (N) filopodia number per 5000 µm^2^ (approximate average neural crest cell area *ex vivo*). Leading edge protrusions were included in cell circularity measurements. DMSO: n=59 cells; 2h BIO: n=35 cells; 2h CHIR99021: n=40 cells analysed, over 3 independent experiments. **** p < 0.0001, *** p < 0.001, ns non-significant, one-way ANOVA, Tukey’s multiple comparisons test. (O-Q) Abi1 immunostaining of fixed migratory cranial neural crest cells, cultured from E8.5 *Wnt1*::cre; *LifeAct-EGFP* embryos, treated for 2 hours with (O) DMSO control, (P) 0.5 µM BIO or (Q) 1 µM CHIR99021. (R) Quantification of the percentage migratory neural crest cells with a positive (white arrow), compromised (open arrowhead), or negative Abi1-positive lamellipodia at their leading edge. Data presented as mean ± SEM. DMSO: n=42 cells; 2h BIO: n=40 cells; 2h CHIR99021: n=39 cells analysed, over 2 independent experiments. *** p < 0.001, * p < 0.05, chi-squared test, scale bar 20 µm. (S-U) Lpd immunostaining of fixed migratory cranial neural crest cells, cultured from CD1 WT E8.5 embryos, treated for 2 hours with (S) DMSO control, (T) 0.5 µM BIO or (U) 1 µM CHIR99021. Scale bar 20 µm. (V) Quantification of the percentage migratory neural crest cells with a positive (white arrow), compromised (open arrowhead), or negative Lpd-positive lamellipodia at their leading edge, following 2 hour treatment with DMSO, BIO or CHIR99021. Data presented as mean ± SEM. DMSO control: n=78 cells; 2h BIO: n=83 cells; 2h CHIR99021: n=60 cells analysed, over 3 independent experiments. **** p < 0.0001, chi-squared test. For Lpd immunostaining following 24 hour GSK3 inhibition, see (Supplementary Figure 1).

We found that migratory neural crest cells treated with GSK3 inhibitors (BIO or CHIR99021) showed reduced speed (MTS) and area explored (Mean Squared Displacement, MSD), compared to DMSO controls (Figure 2G-H). To accurately quantify cell persistence, we used direction autocorrelation (Figure 2I), a measure of how the angle of displacement vectors correlate with themselves^34^. Treatment with the GSK3 inhibitor, CHIR99021, significantly reduced the direction autocorrelation of neural crest cells, compared to DMSO controls (Figure 2I). Altogether, our data suggests that GSK3 promotes neural crest cell migration speed and persistence.

We then asked whether GSK3 activity is required for lamellipodia formation consistent with the previously observed changes in Lpd localisation^13^. We aimed to define the protrusion dynamics of *Wnt1*::cre-expressing neural crest cells by LifeAct-EGFP time-lapse imaging of F-actin dynamics in migratory cells 36 hours after dissection (1 frame/5 sec, 10 min movie) (Figure 2J-L). Neural crest explants were treated with DMSO vehicle control or the GSK3 inhibitors, BIO or CHIR99032, 2 hours prior to imaging. This allowed us to capture the immediate consequences of GSK3 inhibition on lamellipodial dynamics. Wildtype neural crest cells (treated with DMSO as control) showed highly dynamic lamellipodia, presented as colour-coded time-lapse projections, whereby the colour corresponds to frame number (frame 1: red, frame 121: magenta) (Stills: Figure 2J, Supplementary Movie 5). In contrast, neural crest cells treated with BIO or CHIR99021 did not display a lamellipodium (Figure 2K-L) and showed significantly reduced average cell circularity and increased number of filopodia per 5000 μm^2^ average cell area, compared to DMSO controls (Figure 2M-N, Supplementary Movie 5). In slight variation, neural crest cells treated with CHIR99021 had highly dynamic peripheral membrane ruffles which frequently converted into filopodia (Figure 2L, Supplementary Movie 5).

In addition to the actin-rich membrane protrusions at the cell leading edge, high levels of LifeAct-EGFP were also associated with rearward stress fibres in DMSO control neural crest cells (Figure 2K, white arrowhead). In contrast, both BIO- and CHIR99021-treated neural crest cells displayed high intensity F-actin stress fibres extending from the cell rear to the leading edge on all sides of the cells (with the cells appearing less polarised) (Figure 2K-L).

To verify that GSK3 is indeed required for lamellipodia formation, we used immunofluorescence labelling for two key lamellipodial markers, Lpd and Abi1. Abi1 is a component of the Scar/WAVE complex which is required for branched actin nucleation and lamellipodia formation, whilst Lpd acts upstream of the Scar/WAVE complex, directly interacting with Abi1^10,16^. In agreement with other cell types^15,35^, continuous Abi1 and Lpd staining was seen at the edge of the lamellipodium in approximately 80% of DMSO-treated neural crest cells (Figure 2O,S). In contrast, following 2 hour GSK3 inhibitor treatment, only 30-50% of neural crest cells displayed an Abi1-positive lamellipodium (Figure 2R). Instead, cells had punctate Abi1 localisation at the tips of filopodia and/or small remaining areas of Abi1 positive lamellipodia (Figure 2P-Q, quantified in 2R and defined as “compromised”). Similarly, Lpd only displayed continuous leading edge localisation in 40-45% of cells upon GSK3 inhibition, with a concurrent increase in filopodia numbers whose tips were positive for Lpd (Figure 2T-U, quantified in 2V). Furthermore, 24 hour GSK3 inhibition showed similar albeit more severe phenotypes with a loss of lamellipodia and concurrent Lpd localisation (Supplementary Figure 1). This raises the intriguing possibility that GSK3 may regulate lamellipodia formation via Lpd.

### Lamellipodin is required for lamellipodia formation during mouse neural crest migration

We then wanted to define specific roles for *Lpd* in mouse neural crest migration. To do so, we generated neural crest explant cultures from wildtype (*Wnt1*::cre; *Lpd*^+/+^), *Lpd* heterozygous (*Wnt1*::cre; *Lpd*^+/fl^), and homozygous (*Wnt1*::cre; *Lpd*^fl/fl^) knockout mouse embryos (Figure 3A-C, Supplementary Movie 6). We confirmed *Lpd* deletion by immunostaining for Lpd protein in wildtype, heterozygous and homozygous knockout cultures two days following dissection (Supplementary Figure 2). We found a significant reduction in the speed and persistence of *Lpd* knockout cells compared to controls (Figure 3A-F) suggesting that inhibition of GSK3 and knockout of *Lpd* phenocopy each other and thus that they may act in the same pathway.

**Figure 3.**
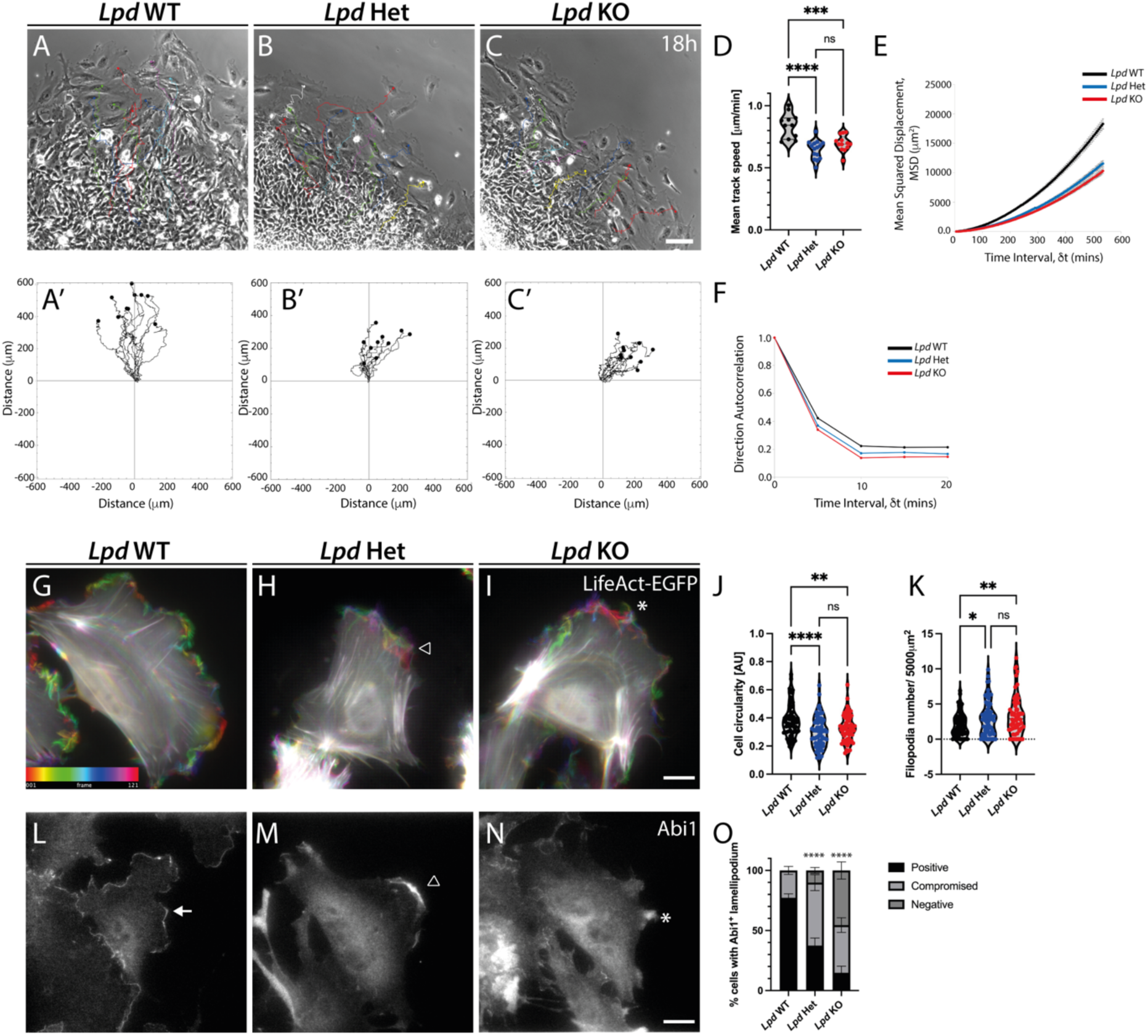
Genetic deletion of *Lpd* prevents lamellipodia formation and reduces the overall migration efficiency of cranial neural crest cells *ex vivo*. **(A-C)** Representative stills from 10x magnification phase-contrast time-lapse imaging of E8.5 **(A)** *Wnt1*::cre; *Lpd*^+/+^, **(B)** *Wnt1*::cre; *Lpd*^+/fl^, and **(C)** *Wnt1*::cre; *Lpd*^fl/fl^ neural crest explant cultures, imaged for 18 hours (1 frame/ 5 min) (see Supplementary Movie 6). Scale bar 100 µm. **(A’-C’)** Trajectory plots of 10 migratory neural crest cells tracked through time-lapse imaging for **(A-C)**. **(D-F)** Quantification of **(D)** Mean Track Speed (MTS), **(E)** Mean Squared Displacement (MSD), and **(F)** direction autocorrelation of migratory neural crest cells. For MTS calculations, each dot represents the mean speed of 10 neural crest cells imaged from the same explant. Lpd WT: n=11 explants, 110 cells tracked; Lpd Het: n=15 explants, 150 cells tracked; Lpd KO: n=13 explants, 130 cells tracked, over 4 independent experiments. **** p < 0.0001, *** p < 0.001, * p < 0.05, ns non-significant, one-way ANOVA, Tukey’s multiple comparisons test. For direction autocorrelation measures, ∂t = 1, TR = 4 (20 min); see Materials & Methods for details. **(G-I)** Colour-coded time projection stills of 60x magnification time-lapse imaging (10 min movies, 1 frame/ 5 seconds) of migratory neural crest cells, cultured from E8.5 **(G)** *Wnt1*::cre; *Lpd*^+/+^; *LifeAct-EGFP*, **(H)** *Wnt1*::cre; *Lpd*^+/fl^; *LifeAct-EGFP*, and **(I)** *Wnt1*::cre; *Lpd*^fl/f^; *LifeAct-EGFP* embryos (see Supplementary Movie 7). Open arrowhead: membrane ruffles; asterisk (*): dynamic filopodia. Scale bar 20 µm. **(J-K)** Quantification of neural crest **(J)** cell circularity and **(K)** filopodia number per 5000 µm^2^ (approximate average neural crest cell area *ex vivo*). Leading edge protrusions were included in cell circularity measurements. Lpd WT: n=68 cells; Lpd Het: n=70 cells; Lpd KO: n=59 cells, analysed over 3 independent experiments. **** p < 0.0001, ** p < 0.01, * p < 0.05, ns non-significant, one-way ANOVA, Tukey’s multiple comparisons test. **(L-N)** Abi1 immunostaining of fixed migratory neural crest cells, cultured from **(L)** *Wnt1*::cre; *Lpd*^+/+^; *LifeAct-EGFP*, **(M)** *Wnt1*::cre; *Lpd*^+/fl^; *LifeAct-EGFP*, and **(N)** *Wnt1*::cre; *Lpd*^fl/fl^; *LifeAct-EGFP* E8.5 embryos. Scale bar 20 µm. **(O)** Quantification of the percentage migratory neural crest cells with a continuous Abi1-positive lamellipodium (positive), a discontinuous Abi1-labelled lamellipodium (compromised) or absent Abi1-labelled lamellipodium (negative). Data presented as mean ± SEM. Lpd WT: n=48 cells; Lpd Het: n=97 cells; Lpd KO: n=71 cells, analysed over 3 independent experiments. **** p < 0.0001, chi-squared test.

Time-lapse imaging of actin dynamics was then completed in *Lpd* wildtype, heterozygous and homozygous knockout neural crest cells, 36 hours after dissection (1 frame/5 sec, 10 min movie) (Stills: Figure 3G-I). Wildtype neural crest cells presented highly dynamic lamellipodial protrusions, presented as colour-coded time-lapse projections, whereby the colour corresponds to frame number (frame 1: red, frame 121: magenta) (Stills: Figure 3G, Supplementary Movie 7). In contrast, neural crest cells with heterozygous expression of *Lpd* displayed highly unstable membrane ruffles at the cell leading edge (Figure 3H, arrowhead, Supplementary Movie 7), with an overall reduced cell circularity (Figure 3J). More drastically, *Lpd* conditional knockout neural crest cells lacked a filopodial protrusions (Figure 3I-K, asterisk) which, were highly dynamic and after initial outward extension, flipped backward into the lamella (Supplementary Movie 7).

To unambiguously detect lamellipodia, we stained wildtype, *Lpd* heterozygous and *Lpd* homozygous knockout neural crest explants for the lamellipodial marker Abi1 (Figure 3L-N). In wildtype neural crest cells, approximately 80% of cells had continuous Abi1 immunostaining, indicative of lamellipodia formation (Figure 3L, arrow, 3O). In *Lpd* heterozygous neural crest cells, 50% displayed a compromised lamellipodium in which protrusions appeared as unstable ruffles (Figure 3M, arrowhead, 3O). Most *Lpd* knockout cells lacked an Abi1-positive lamellipodium, with only 15% retaining small areas of lamellipodia (Figure 3N, asterisk, 3O). These data identify a novel requirement for *Lpd* in lamellipodia formation in primary mouse cranial neural crest cells. This change in actin dynamics, loss of lamellipodia, and induction of filopodia upon loss of *Lpd* is reminiscent of phenotypes reported for cells lacking the Arp2/3 complex, an essential nucleator of lamellipodial branched actin^29^. This indicates that Lpd may be functioning upstream of Scar/WAVE-Arp2/3 complexes^16^ in neural crest cells.

### Lamellipodin controls actin-based protrusions in mouse neural crest cells in vivo

To determine whether Lpd regulates actin-based protrusions *in vivo*, we generated mice carrying neural crest-specific expression of the membrane-EGFP reporter with or without conditional *Lpd* knockout^16,30,32^. *Lpd* knockout in the neural crest lineage significantly reduced the number of *Lpd* knockout pups born at early postnatal stage, P0 (Supplementary Figure 3). Moreover, a significant difference was seen between the expected and observed number of *Lpd* knockout embryos at the early embryonic timepoint, E8.5 (Supplementary Figure 3). However, we did not observe gross morphological differences when comparing wildtype, *Lpd* heterozygous (*Wnt1*::cre; *Lpd*^+/fl^; *Rosa26R^mtmg^)* and *Lpd* homozygous knockout (*Wnt1*::cre; *Lpd*^fl/fl^; *Rosa26R^mtmg^) e*mbryos at E8.5 or E9.5 (Figure 4A-C, G-H).

**Figure 4.**
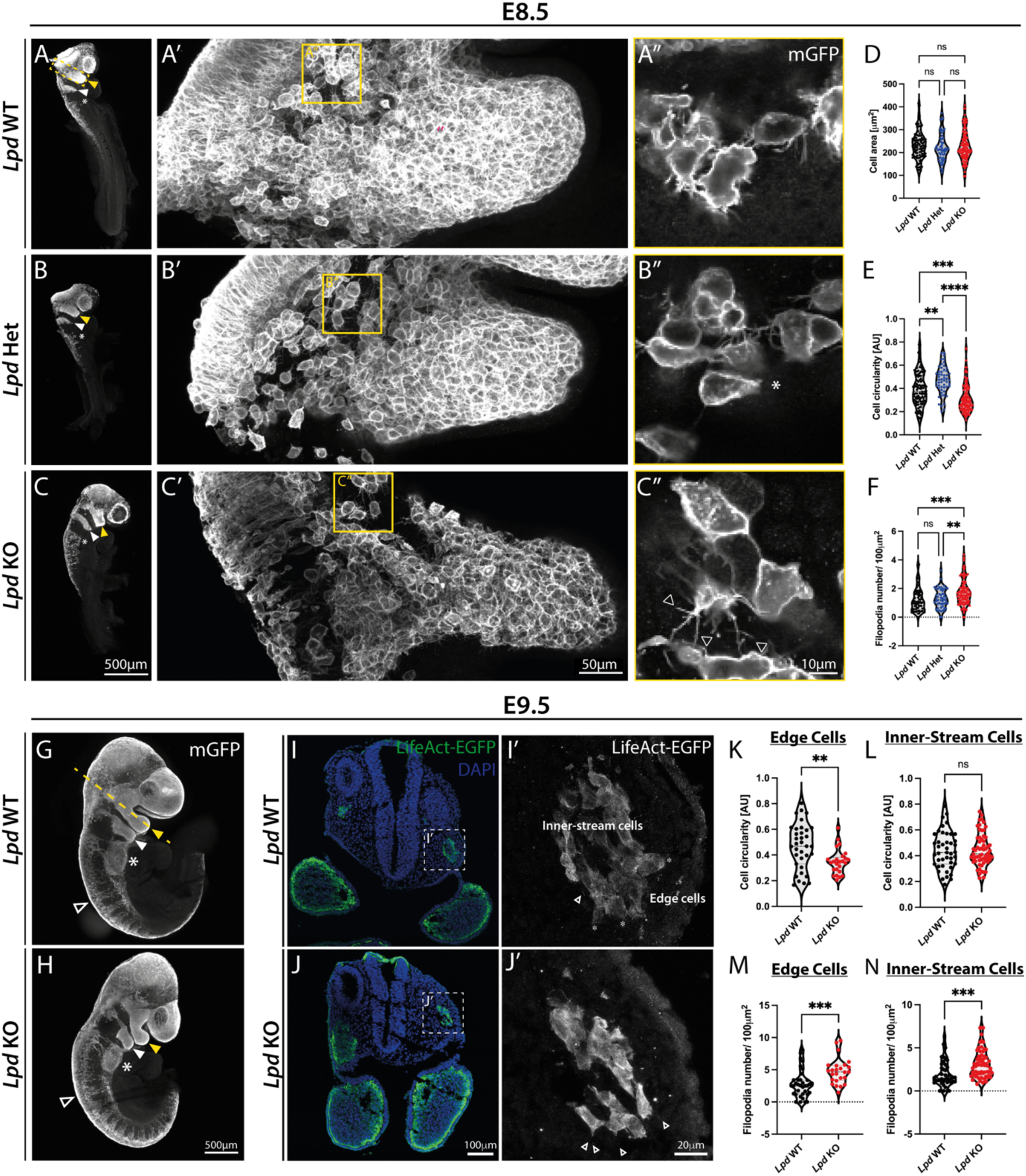
Genetic deletion of *Lpd* increases the number of filopodia protrusions in cranial neural crest cells *in vivo*. **(A-C)** Maximum intensity projections of E8.75 (6-10 somite stage) **(A)** *Wnt1*::cre; *Lpd*^+/+^; *Rosa26^MTMG^,* **(B)** *Wnt1*::cre; *Lpd*^+/fl^; *Rosa26^MTMG^* and **(C)** *Wnt1*::cre; *Lpd*^fl/fl^; *Rosa26^MTMG^* embryos, whose *Wnt1*::cre-driven membrane-GFP expression has been retrieved using an anti-GFP antibody. Scale bar 500 µm. Dashed box indicates imaging region of **(A’-C’)**. Pharyngeal arch-1 (PA1), yellow arrowhead), PA2 (white of the neural crest stream migrating towards PA1 (70 µm z-stacks). Scale bar 50 µm. **(A’’-C’’)** Single z-stack optical slices from neuroepithelium-adjacent locations **(A’-C’)**. Scale bar: 10 µm. **(D-F)** Quantification of **(D)** cell area, **(E)** cell circularity and **(F)** filopodia number per 100 µm^2^. Each dot represents one cell (Lpd WT: N = 70, Lpd Het: N = 53, Lpd KO: N = 109), from at least 2 embryos per genotype over 3 independent experiments. ** p < 0.01, *** p < 0.001, **** p <0.0001, ns non-significant, one-way ANOVA and Tukey’s multiple comparison’s test. **(G-H)** Maximum intensity projection images of **(G)** *Wnt1*::cre; *Lpd*^+/+^; *Rosa26^MTMG^* and **(H)** *Wnt1*::cre; *Lpd*^fl/fl^; *Rosa26^MTMG^* E9.5 embryos. Scale bar 500 µm. PA1: yellow arrowhead, PA2 (white arrowhead), cardiac/ vagal regions (*), trunk neural crest streams (open arrowhead). **(I-J)** Maximum projections of transverse sections through the head of E9.5 **(I)** *Wnt1*::cre; *Lpd*^+/+^; *LifeAct-EGFP* or **(J)** *Wnt1*::cre; *Lpd*^fl/fl^; *LifeAct-EGFP* embryos. *Wnt1*::cre-driven LifeAct-EGFP (green) marks neural crest contributions, and DAPI (blue) marks nuclei. Scale bar 100 µm. **(I’-J’)** 60x magnification (2x zoom) maximum projections of **(I)** and **(J)**, indicated by a white dotted box. Scale bar 20µm. **(K-L)** Quantification of neural crest cell circularity, at **(K)** the edge of the neural crest stream or **(L)** within the stream. Leading edge protrusions including filopodia were included in the cell circularity measurements. **(M-N)** Quantification of filopodial protrusions/100 µm^2^, at **(M)** the edge of the stream or **(N)** within the stream. Each dot represents one cell (edge cells: Lpd WT: N = 35, Lpd KO: N = 25; inner-stream cells: Lpd WT: N = 43, Lpd KO: N = 60) analysed over 3 independent experiments. ** p < 0.01, *** p < 0.001, ns non-significant, unpaired t test.

Nevertheless, at somite stage 6-10 (E8.75), we observed a significant increase in the number of filopodial protrusions in homozygous *Lpd* knockout neural crest cells *in vivo* (Figure 4C”) compared to wildtype (Figure 4A”, quantified in Figure 4F). This suggest that Lpd is also required *in vivo* for lamellipodia formation and in its absence lamellipodial protrusions are replaced by filopodia. Furthermore, despite similar size (area) of neural crest cells in all genotypes (Figure 4B”, D), average cell circularity was increased in *Lpd* heterozygous cells and decreased in *Lpd* homozygous mutant cells (Figure 4C”, E).

By 24-26 somite stage (E9.5) neural crest cells have filled the first and second pharyngeal arches (yellow and white arrowheads), cardiac/vagal neural crest cells (asterisk) have migrated more ventrally, and more posterior trunk crest streams are migrating (e.g. at somite 16) (open arrowhead) (Figure 4G). To examine whether the role of Lpd in regulating cranial neural crest cell protrusions was conserved at E9.5 *in vivo*, we examined the cranial neural crest streams migrating away from the dorsal neural tube and towards pharyngeal arch 1 in transverse cryo-sections through the head of E9.5 embryos (Figure 4I-J). We examined cells at the edge of the migratory stream, and from within the inner neural crest stream (Figure 4I’-J’). For both locations we found that *Lpd* homozygous knockouts were more elongated compared to wildtype controls (Figure 4I’-J’, K-L), and displayed an increased number of filopodia/100μm^2^ (Figure 4M-N). Together, these data support a role for Lpd in counteracting filopodia formation (whilst potentially supporting lamellipodia formation) in cranial neural crest cells *in vivo*.

### Lpd is a substrate of GSK3 kinase

Since we observed that inhibition of GSK3 activity and loss of *Lpd* phenocopy each other, we hypothesised that GSK3 might directly phosphorylate Lpd to control its function at the leading edge of cells. To test this, we expressed an Lpd-EGFP fusion protein in HEK293FT cells in the presence or absence of a constitutively active HA-tagged GSK3β (HA-GSK3β-DA). Following immunoprecipitation (IP) of EGFP-tagged Lpd, we found that HA-GSK3β-DA weakly associated with Lpd-EGFP (Figure 5A, lane 6). Co-expression of HA-GSK3β-DA with Lpd-EGFP also increased overall serine-threonine phosphorylation of Lpd, but reduced Lpd threonine phosphorylation (Figure 5A, lane 6), suggesting that GSK3β may preferentially phosphorylate Lpd on serine residues.

**Figure 5.**
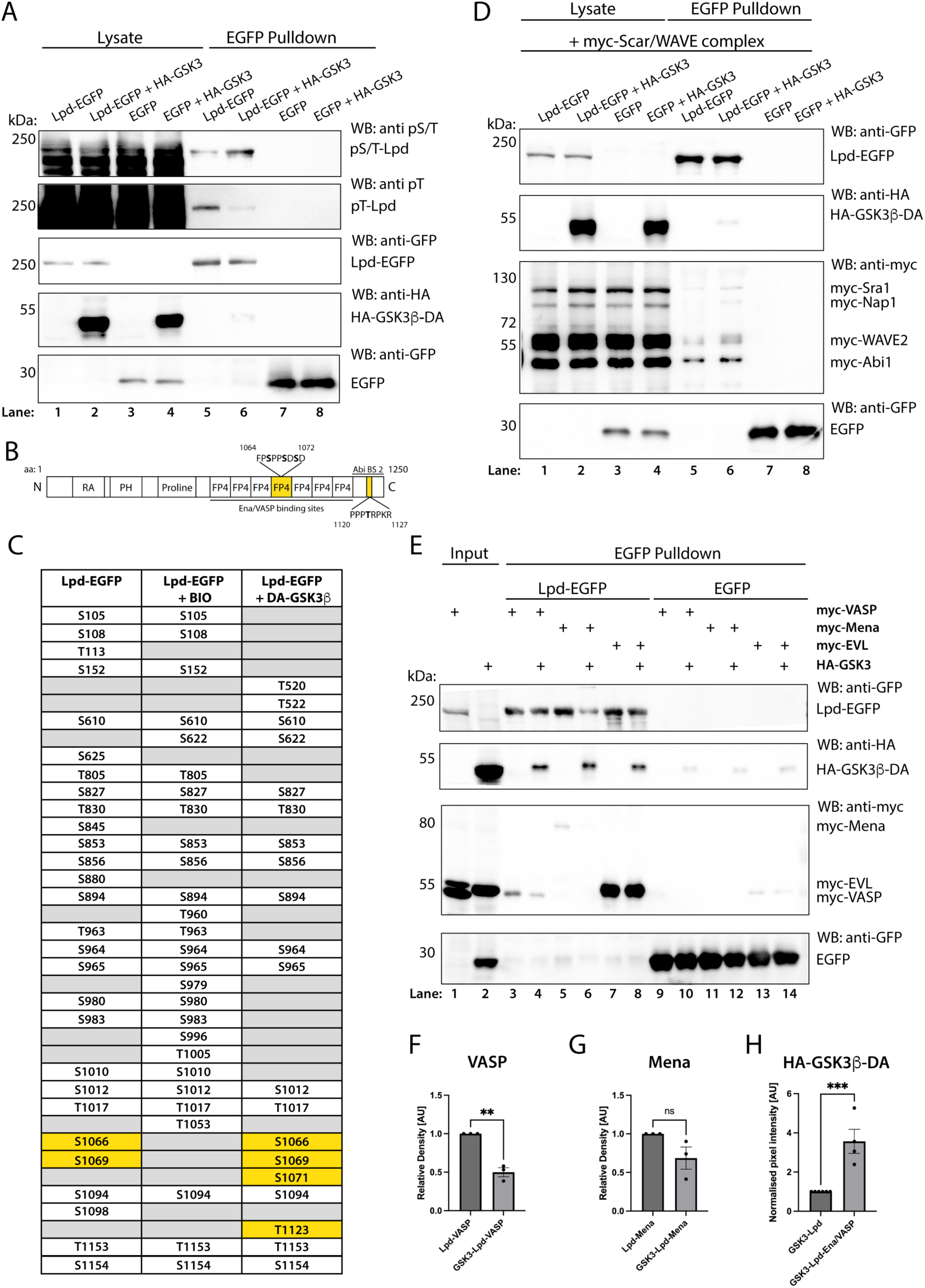
GSK3 phosphorylates Lpd to increase Lpd interaction with Scar/WAVE, and reduces interaction with Ena/VASP proteins, VASP and Mena. **(A)** HA-tagged, dominant-active (DA) GSK3b (DA-GSK3b-HA) co-immunoprecipitation with Lpd-EGFP in HEK293FT cells. EGFP-Trap pulldowns were performed from cell lysates followed by western blotting and probing with anti-EGFP, anti-HA, anti-phospho-serine/threonine (pS/T) and anti-phospho-threonine (pT) antibodies. Blots are representative of 3 independent experiments. **(B)** Schematic representation of Lpd protein domain structure. RA: Ras-association domain, PH: Pleckstrin homology domain, Proline: proline-rich region, FP4: FPPPP motif (Ena/VASP binding sites). GSK3b phosphorylates Lpd at S1066, S1069 and S1071 within Lpd FP4-4 motif, an Ena/VASP binding site, and T1123 within Abi1 binding site 2, marked in bold (yellow boxes). **(C)** Summary table of differentially phosphorylated serine-threonine residues on Lpd in the presence or absence of GSK3b. Column 1: Phosphorylated Lpd S/T residues with Lpd-EGFP overexpression. Column 2: Phosphorylated Lpd S/T residues with inhibition of GSK3 and Lpd-EGFP overexpression. Column 3: Phosphorylated Lpd S/T residues with co-expression of DA-GSK3b-HA and Lpd-EGFP. Yellow boxes: GSK3 phosphorylation sites of interest, grey boxes: non-phosphorylated residues. **(D)** DA-GSK3b-HA increases the co-immunoprecipitation of myc-tagged Scar/WAVE complex components with Lpd-EGFP in HEK293FT cells. Blots were probed for anti-myc, anti-HA and anti-EGFP, and are representative of 3 independent experiments. **(E)** DA-GSK3b-HA reduces the co-immunoprecipitation of myc-tagged Ena/VASP proteins, myc-VASP and myc-Mena, with Lpd-EGFP in HEK293FT cells. Blots were probed for anti-myc, anti-HA and anti-EGFP, and are representative of 3 independent experiments. For the lysate control blot, see (Supplementary Figure 4). **(F-H)** Quantification of normalised pixel intensity of **(F)** myc-VASP, **(G)** myc-Mena and **(H)** DA-GSK3b-HA, normalised to Lpd-EGFP co-immunoprecipitation band intensity. *** p < 0.001, ** p < 0.01, ns non-significant, unpaired t-test. Each dot represents 1 independent experiment (n=3).

To map the GSK3 phosphorylation sites on Lpd, we expressed the Lpd-EGFP fusion protein in HEK293FT cells with or without treatment with the GSK3 inhibitor, BIO, or co-expression of Lpd-EGFP with HA-GSK3β-DA. After immunoprecipitation of Lpd-EGFP, tandem mass spectrometry was performed. By comparing these loss- and gain-of-function scenarios, we observed GSK3-dependent changes in multiple Ser/Thr phosphorylation sites throughout Lpd (Figure 5B-C). Lpd is known to interact with the N-terminal EVH1 domain of Ena/VASP proteins via 7 proline-rich motifs characterised by a core motif composed of phenylalanine followed by four prolines which is flanked by acidic amino acids (FP4 motifs) (Figure 5B)^15^. Three GSK3 phosphorylation sites in Lpd were therefore of particular interest: GSK3β overexpression was associated with phosphorylation at three serine residues (S1066, S1069, S1071) in the C-terminus, specifically within the fourth Lpd FP4 motif (FP4-4) (Figure 5C, right) two of which were no longer phosphorylated upon GSK3 inhibition (Figure 5C, Ser-1066, Ser-1069).

In addition, GSK3 was also associated with phosphorylation at T1123, a threonine within the second Abi1 binding site on Lpd^36^ (Figure 5C, right). These data suggest that Lpd is a novel substrate of GSK3, and that GSK3 phosphorylation of Lpd may regulate its interactions with Ena/VASP proteins and/or the Scar/WAVE complex.

### GSK3 promotes Lpd interactions with the Scar/WAVE complex and reduces interactions with Ena/VASP proteins

As noted, a key function of Lpd is to act as a scaffold for the major actin effectors, Scar/Wave-Arp2/3 complexes or Ena/VASP proteins. Based on the mass spectrometry data, we proposed that GSK3-dependent phosphorylations alter Lpd interactions with these partner proteins. First, we tested the Lpd-Scar/WAVE interaction by co-expressing Lpd-EGFP and Myc-tagged Scar/WAVE complex components (Sra1, Nap1, Scar/WAVE2, Abi1, HSPC300) with or without HA-GSK3β-DA (Figure 5D). Co-immunoprecipitation of Lpd with the Scar/WAVE complex was increased in the presence of dominant active GSK3β (Figure 5D, lane 6).

The analogous Ena/VASP interaction experiment was performed with Lpd-EGFP, Myc-tagged Ena/VASP proteins (VASP, Mena or EVL) and HA-GSK3β-DA (Figure 5E, EGFP pulldown). The full protein lysate control blot can be found in (Supplementary Figure 4). When not co-expressed with GSK3β, Lpd-EGFP showed the strongest interaction with the Ena/VASP protein, EVL (Figure 5E, lane 7), and a weaker interaction with Mena and VASP (Figure 5E, lanes 3, 5). Surprisingly, in contrast to the Lpd-Scar/WAVE complex interaction, co-expression of HA-GSK3β-DA reduced the interaction of Lpd with VASP and Mena, but not with EVL (Figure 5E, lanes 4,6, quantified in Figure 5F-G). We also found that the co-expression of Lpd-EGFP with Myc-Ena/VASP led to an increase in the amount of HA-GSK3β-DA associated with Lpd-EGFP in the IP (compare Figure 5D lane 6 with Figure 5E, lanes 4, 6, 8, and quantified in Figure 5H), suggesting the formation of a stable complex between Lpd, GSK3 and Ena/VASP proteins. Together this suggests that GSK3 acts to promote Lpd interactions with the Scar/WAVE complex but reduces Lpd interactions with the Ena/VASP proteins, VASP and Mena.

### GSK3 and Lpd promote Ena/VASP localisation to the leading edge at the expense of focal adhesions in mouse neural crest cells

We then turned back to the *Lpd* genetic mutants to determine whether endogenous Lpd is required for Ena/VASP protein localisation during neural crest cell migration. Both Mena and VASP (Figure 6A, L’, Supplementary Figure 6-7) are expressed in migratory cranial neural crest cells, and double *Mena/VASP* knockout mice display neurulation and craniofacial defects, indicative of an essential function in the mouse neural crest^37,38^. We focused here on Mena localisation at the very edge of the lamellipodium and in focal adhesions. During lamellipodia formation, adhesions occur underneath the newly formed protrusion, and focal adhesion maturation must occur before cell rear contraction in order to sustain efficient contractile forces^39–41^. As expected from other cell types^42^, Mena predominantly localises to mature adhesions at the cell rear and to the lamellipodia in mouse neural crest cells (Figure 6A-A’, L).

**Figure 6.**
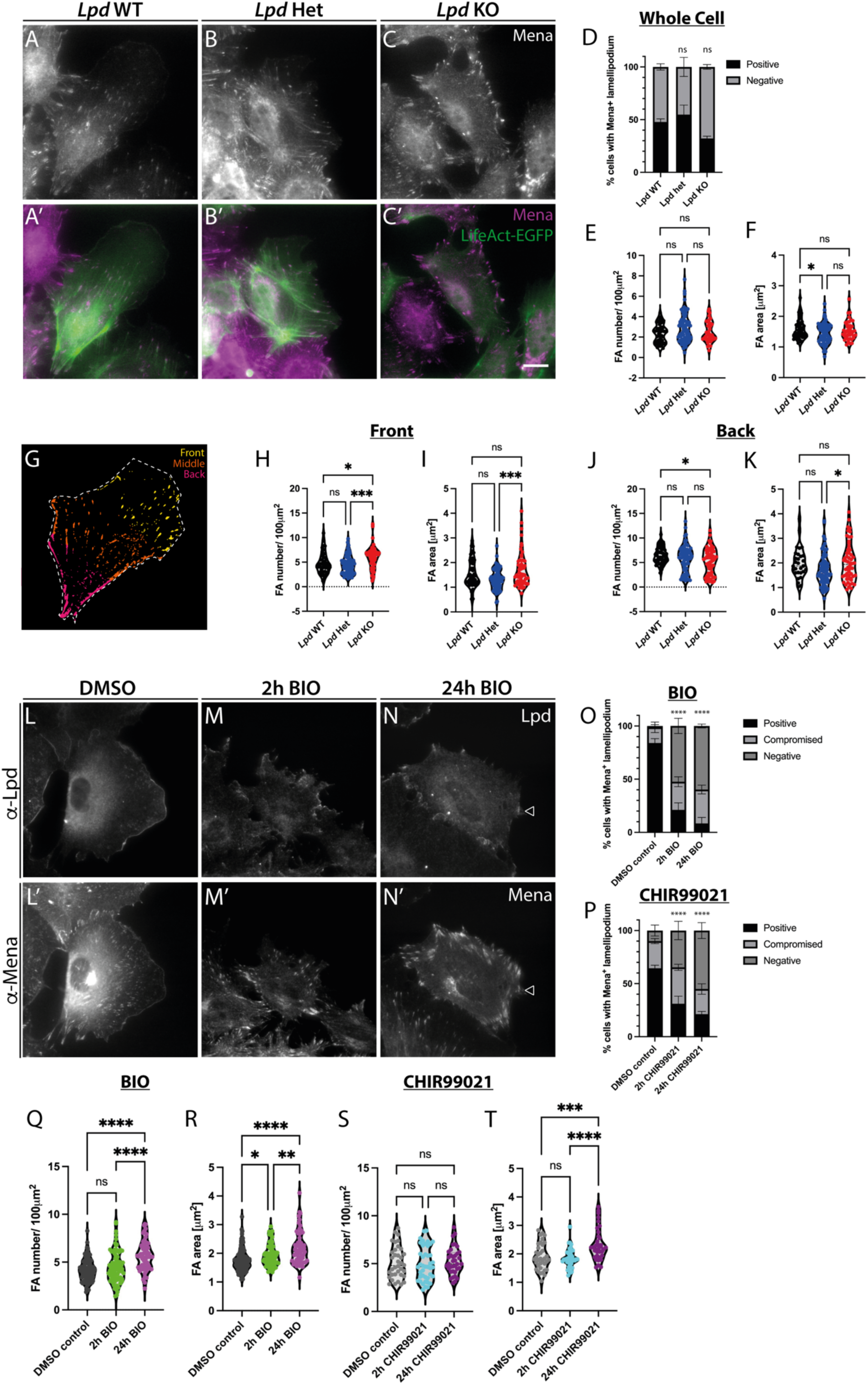
*Lpd* deletion and GSK3 inhibition mislocalise Ena/VASP proteins to mature focal adhesions at the front of mouse cranial neural crest cells. (A-C) Mena immunostaining of fixed migratory cranial neural crest cells, cultured from E8.5 (A) *Wnt1*::cre; *Lpd*^+/+^; *LifeAct-EGFP*, (B) *Wnt1*::cre; *Lpd*^+/fl^; *LifeAct-EGFP*, and (C) *Wnt1*::cre; *Lpd*^fl/fl^; *LifeAct-EGFP* embryos. (A’-C’) Merged images of a-Mena (magenta) and LifeAct-EGFP (green). Scale bar 20 µm. (D) Quantification of the percentage neural crest cells with Mena localisation to the lamellipodium. Data presented as mean ± SEM, and analysed using a chi-squared test. (E-F) Quantification of the average (E) Mena-positive focal adhesion number/ 100 µm^2^, and (F) focal adhesion area in *Lpd* wild-type, conditional heterozygous and homozygous knockout cranial neural crest cells. (G) Segmentation strategy used to sub-classify Mena-positive focal adhesions according to their localisation to the front, middle and back thirds of cells. (H-I) Quantification of (H) average Mena-positive focal adhesion number/ 100 µm^2^ and (I) average focal adhesion area within the front third of cells. (J-K) Quantification of (J) average Mena-positive focal adhesion number/ 100 µm^2^ and (K) average focal adhesion area within the back third of cells. Each dot represents one cell. *** p < 0.001, * p<0.05, ns non-significant, one-way ANOVA, Tukey’s multiple comparisons test. Lpd WT: N = 48, Lpd Het: N = 124, Lpd KO: N = 72, over 3 independent experiments. (L-N’) Lpd (L-N) and Mena (L’-N’) co-immunostaining of fixed migratory cranial neural crest cells, cultured from E8.5 WT embryos, treated with (L-L’) DMSO, or the GSK3 inhibitor BIO, for (M-M’) 2 hours or (N-N’) 24 hours prior to fixation. For immunofluorescence images of Lpd-Mena treated with CHIR99021, see (Supplementary Figure 5). (O-P) Quantification of the percentage neural crest cells with Mena localisation to the lamellipodium. Data presented as mean ± SEM. **** p < 0.0001, analysed using a chi-squared test (n=3). (Q-T) Quantification of the average Mena-positive (Q,S) focal adhesion number/ 100 µm^2^ and (R,T) focal adhesion area in the whole cell of DMSO control, BIO-treated (Q-R) or CHIR99021-treated (S-T) cranial neural crest cells, 2 or 24 hours prior to fixation. Each dot represents one cell, from 3 independent experiments. **** p < 0.0001, ** p < 0.01, * p<0.05, ns non-significant, one-way ANOVA, Tukey’s multiple comparisons test. DMSO: N = 99, 2h BIO: N = 79, 24h BIO: N = 61, 2h CHIR99021: N = 64, 24h CHIR99021: N = 37 cells.

When we quantified the localisation to lamellipodia, we found that 80% of wildtype neural crest cells display lamellipodia but Mena only localised to 50% of lamellipodia (Figure 6A, D). This suggests that, at least in neural crest cells, Mena is not a constitutive component of lamellipodia. As noted above, in *Lpd* heterozygous and homozygous knockout cells, most migratory cells did not have a lamellipodium (Figure 3N-P). However, 55% of heterozygous and 30% of homozygous Lpd knockout neural crest cells displayed small ruffles positive for Mena instead of a lamellipodium (Figure 6B-C’,D). This lack of lamellipodia meant that we were unable to definitively conclude whether Lpd was required for leading edge localisation of Mena.

We then assessed the requirements for Lpd in the recruitment of Mena to focal adhesions. We used antibody staining to analyse endogenous Mena localisation at the whole cell level (Figure 6E-F) and within the front, middle and back thirds of the cell (Figure 6G-K). At the whole cell level, no significant difference was seen in the average number of Mena-positive focal adhesions per 100 μm^2^ nor average Mena-positive focal adhesion area between *Lpd* wildtype and *Lpd* homozygous knockout neural crest cells (Figure 6E-F). However, within the cell front, *Lpd* knockout neural crest cells had a significantly increased number of Mena-positive mature focal adhesions, compared to controls and *Lpd* heterozygous cells (Figure 6H). Conversely, within the back third of cells, there was a significant reduction in the number of Mena-positive focal adhesions (Figure 6J). Together, this suggests that genetic loss of *Lpd* shifts Mena-positive mature focal adhesions towards the front of mouse cranial neural crest cells. This may be a consequence of loss of more quickly.

We then checked whether GSK3-dependent phosphorylation affected Lpd-Mena co-localisation. Wildtype explants were treated with DMSO control or GSK3 inhibitors (BIO or CHIR99021) for 2h or 24h prior to fixation. Explants were co-immunostained for Lpd and Mena (Figure 6L-N’). In DMSO controls, Lpd and Mena, as expected, showed high levels of co-localisation at the lamellipodial edge (Figure 6L-L’). Following 2h and 24h BIO treatment, significantly less cells had Mena- and Lpd-positive lamellipodia (Figure 6O-P). Notably, both BIO and CHIR99021 dramatically increased the size of Mena-positive focal adhesions, as well as increased focal adhesion number upon BIO treatment (Figure 6M-N’, Q-T, Supplementary Figure 5).

This effect occurred rapidly, within 2 hours of treatment, and was reversible upon the washout of pharmacological inhibitor (Supplementary Figure 6). Wash-out progressively reversed cell front localisation of Mena-positive mature focal adhesions, which was reinstated by 15h BIO wash-out (Supplementary Figure 6A, C-F). Similarly, re-localisation of Lpd and Mena to the edge of lamellipodial protrusions was seen by 18 hour washout (Supplementary Figure 6B). BIO wash-out was also able to rapidly and reversibly control VASP localisation within cranial neural crest cells (Supplementary Figure 7). These experiments combined suggest that GSK3 phosphorylates Lpd to counteract Ena/VASP recruitment to focal complexes at the leading edge of migratory neural crest cells.

### Lpd prevents focal complex maturation

Recruitment of Ena/VASP proteins to early adhesions is mediated by direct interaction with the Lpd-related protein, RIAM^43^. Ena/VASP proteins then promote focal adhesion maturation and cell spreading^27,28,42–48^. RIAM is required for integrin activation through a direct interaction with talin^43,49–51^. RIAM binds to talin in nascent adhesions, and must be displaced by vinculin for maturation of focal complexes^52^.

Lpd may play a similar role to RIAM in neural crest cells as single cell RNAseq revealed that Lpd but not RIAM is expressed in cranial neural crest cells^53^. Therefore, we examined the distribution of vinculin, to mark focal complexes, and zyxin which only appears after maturation into focal adhesions^54–57^. As expected, vinculin was localised to focal complexes behind the lamellipodial edge and mature adhesions throughout the cell (Figure 7A-A’), whilst zyxin was localised to stress fibres and mature adhesions at the rear of wildtype cells (Figure 7H-H’). Upon *Lpd* deletion, the area but not the number of vinculin-positive adhesions dramatically increased, with altered localisation to the cell periphery (Figure 7C-C’, D-G) suggesting that these represent more mature focal adhesions.

**Figure 7.**
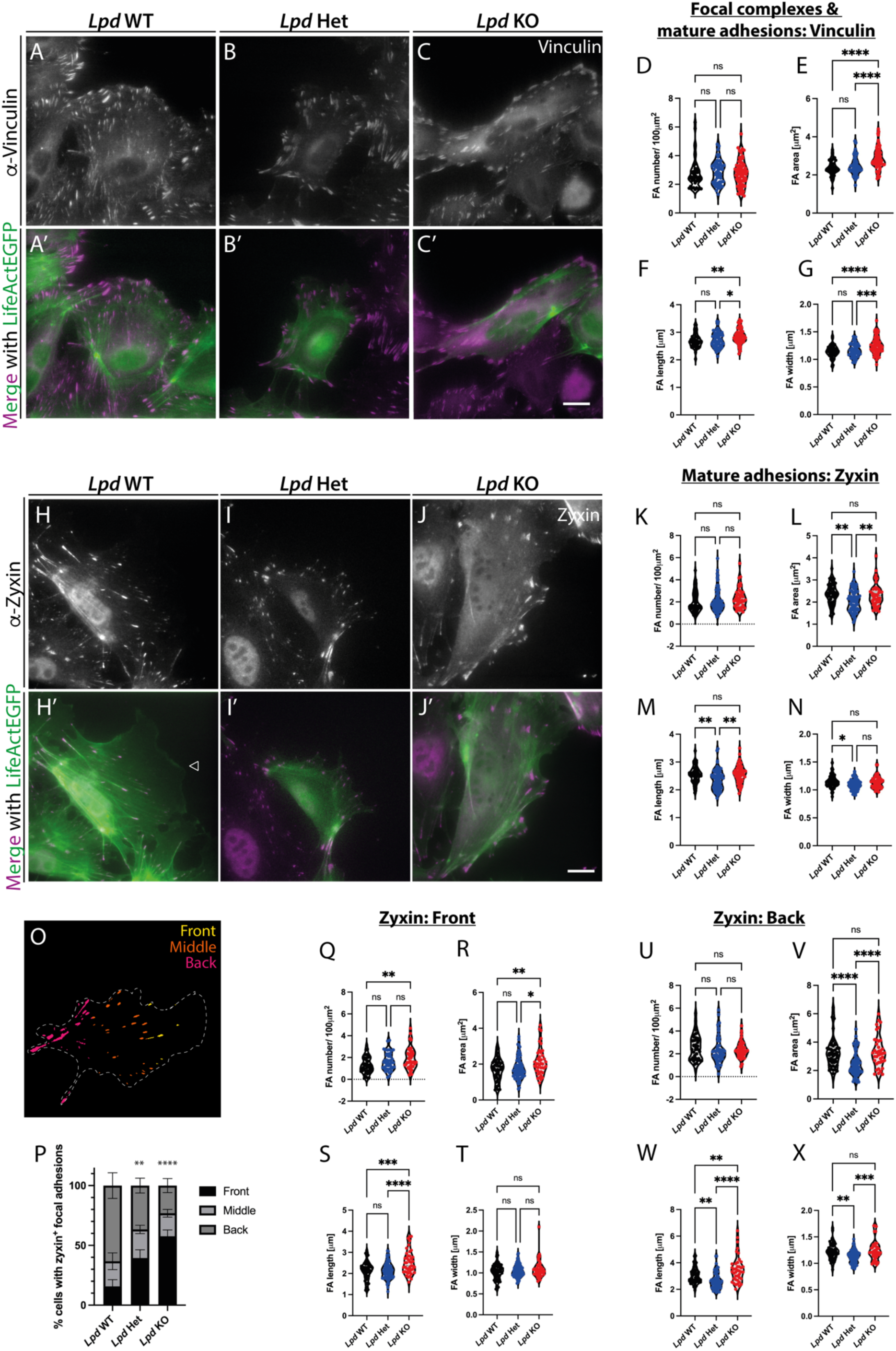
Lpd inhibits nascent adhesion maturation at the leading edge of mouse cranial neural crest cells. **(A-C)** Vinculin immunostaining of fixed migratory cranial neural crest cells, cultured from E8.5 **(A)** *Wnt1*::cre; *Lpd*^+/+^; *LifeAct-EGFP*, **(B)** *Wnt1*::cre; *Lpd*^+/fl^; *LifeAct-EGFP* and **(C)** *Wnt1*::cre; *Lpd*^fl/fl^; *LifeAct-EGFP* embryos. **(A’-C’)** Merged immunostaining of Vinculin (magenta) with LifeAct-EGFP fusion protein (green). Scale bar 20 µm. **(D-G)** Quantification of Vinculin-positive average **(D)** focal adhesion number/ 100 µm^2^, **(E)** focal adhesion area, **(F)** focal adhesion length and **(G)** focal adhesion width in *Lpd* wildtype, *Lpd* heterozygous and homozygous knockout cells. **(H-J)** Zyxin immunostaining of fixed migratory neural crest cells, cultured from E8.5 **(H)** *Wnt1*::cre; *Lpd*^+/+^; *LifeAct-EGFP*, **(I)** *Wnt1*::cre; *Lpd*^+/fl^; *LifeAct-EGFP* and **(J)** *Wnt1*::cre; *Lpd*^fl/fl^; *LifeAct-EGFP* embryos. **(H’-J’)** Merged immunostaining of zyxin (magenta) with LifeAct-EGFP fusion protein (green). Scale bar 20 µm. **(K-N)** Quantification of Zyxin-positive average **(K)** focal adhesion number/ 100 µm^2^, **(L)** focal adhesion area, **(M)** focal adhesion length and **(N)** focal adhesion width in *Lpd* wild-type, *Lpd* heterozygous and homozygous knockout cells. Each dot represents one cell (Lpd WT: N = 42, Lpd Het: N = 52, Lpd KO: N = 73), over at least 3 independent experiments. **(O)** Segmentation strategy used to sub-classify Zyxin-positive focal adhesions according to their localisation to the front, middle and back thirds of neural crest cells. **(P)** Quantification of zyxin-positive focal adhesion localisation to the front, middle or back thirds of *Lpd* wild-type, heterozygous and homozygous knockout neural crest cells. Data presented as mean ± SEM. **** p < 0.0001, ** p < 0.01, analysed using a chi-squared test. **(Q-T)** Quantification of zyxin-positive average **(Q)** focal adhesion number/ 100 µm^2^, **(R)** focal adhesion area, **(S)** focal adhesion length and **(T)** focal adhesion width in the front third of *Lpd* wild-type, heterozygous and homozygous knockout cells. **(U-X)** Quantification of zyxin-positive average **(U)** focal adhesion number/ 100 µm^2^, **(V)** focal adhesion area, **(W)** focal adhesion length and **(X)** focal adhesion width in the back third of cells. Each dot represents one cell (Lpd WT: N = 58, Lpd Het: N = 78, Lpd KO: N = 44), over at least 3 independent experiments. **** p < 00001, *** p < 0.001, ** p < 0.01, * p < 0.05, ns non-significant, one-way ANOVA, Tukey’s multiple comparisons test.

At the whole cell level, the overall numbers and area of zyxin-positive focal adhesions did not change significantly in mutants (Figure 7J-J’, K-N). However, a significant change in the distribution of zyxin-positive adhesions between the front, middle and back third of neural crest cells was apparent (Figure 6O-P). At the front of the cell, both the number and length of zyxin-positive mature focal adhesions was increased in *Lpd* knockout cells (Figure 7Q, S), with the zyxin-positive focal adhesion length also increased at the cell rear (Figure 7W). These data suggest that *Lpd* genetic deletion causes a premature maturation of focal complexes. We therefore propose that Lpd promotes lamellipodia formation through its interaction with the Scar/WAVE complex and by controlling focal complex maturation through its interaction with Mena and VASP.

## DISCUSSION

Here, we report that mouse cranial neural crest cells use lamellipodia and filopodia *in vivo*. By culturing primary neural crest cells, we show that inhibition of GSK3 or deletion of *Lpd* causes defective lamellipodia formation and focal adhesion maturation. Loss of GSK3 and Lpd phenocopy one another, indicating that both are required to promote cranial neural crest migration. We then identify GSK3-dependent phosphorylation of Lpd on many S/T sites, including in an Ena/VASP binding site and a Scar/WAVE complex binding site. GSK3 activity increases Lpd binding to the Scar/WAVE complex, whilst decreasing Lpd-VASP and -Mena interactions. Thus, GSK3 may promote lamellipodia formation through Lpd-Scar/WAVE and inhibit focal adhesion maturation by reducing the recruitment of VASP and Mena to Lpd at these sites.

Neural crest cells are known to have species-specific and neural crest stream-specific cellular behaviours^2,14^. Differences in actin-based protrusions have also been observed, with filopodia predominantly used *in vivo* in the zebrafish neural crest^14,58^, whilst both lamellipodia and filopodia are used in chicken and *Xenopus* cranial neural crest cells *in vivo*^59–61^. However, actin-based protrusions are not well-described in mouse. We previously observed lamellipodia in mouse neural crest cells in culture, and the loss thereof in *GSK3* knockouts^13^. However, the *in vivo* protrusions used in mouse neural crest cell populations are unknown. Here, we provide evidence for the use of lamellipodia in mouse cranial neural crest cells *in vivo*. Notably, lamellipodia are seen in early delaminating neural crest cells, whilst in later streams filopodia predominate (Figure 1). Our data is consistent with the known role of filopodia in sensing and responding to chemo-attractive and durotactic signals, crucial to migration in complex 3D environments^8,12,60, 62–65^, such as the *in vivo* cranial neural crest streams, migration of the embryonic mesoderm wings^66^ and myotube migration through the *Drosophila* testes^67^. These studies highlight the importance of both lamellipodia and filopodia during *in vivo* and developmental mesenchymal migration programmes.

The requirement for Lpd in lamellipodia formation in mouse neural crest cells (Figure 3) is complementary to previous studies in *Lpd* knockdown B16-F1 melanoma cells and *Lpd* conditional knockout mouse embryonic fibroblasts^15,16^. This contrasts with a report using CRISPR-Cas9-driven permanent *Lpd* knockout in B16-F1 melanoma cells which found Lpd to be dispensable for lamellipodia formation^68^, suggesting that the permanent knockout of *Lpd* induces compensatory upregulation of other genes that maintain lamellipodium architecture. *Arp2/3* knockout mouse embryonic fibroblasts and neural crest-derived melanoblasts lacking *Arp2/3* also present an increased number of filopodia at the cell edge^29,69^. These phenotypes are in agreement with our study, which complements previous reports of Lpd-Scar/WAVE interactions promoting *Xenopus* cranial neural crest migration^16^.

A key novelty in our work is the identification of Lpd as a substrate for GSK3 (Figure 5). GSK3 is known to be highly promiscuous, with many potential target substrates^70^. However, very few of these target proteins have been challenged using both loss- and gain-of-function experiments^71,72^. As such, Lpd is now one of only a few functionally validated GSK3 substrates that can directly regulate actin-based motility downstream of Rac1 (Figure 5)^10^. This is in contrast to known GSK3 substrates that regulate mesenchymal migration via microtubule stability downstream of Cdc42^73–77^, as well as focal adhesion dynamics via FAK^78,79^.

At least two distinct tyrosine kinases, c-Src and c-Abl, have been reported to regulate Lpd^17,80^. Phosphorylation of Lpd by c-Src promotes Lpd interaction with the Scar/WAVE complex *in vitro*, whilst c-Abl phosphorylation increases Lpd interactions with both Ena/VASP proteins and the Scar/WAVE complex during 3D cancer cell invasion^17,80^. However, GSK3 is the first serine/threonine kinase shown to phosphorylate Lpd that acts as a dual regulator, where activation of one pathway (Scar/WAVE) is favoured over the other (Ena/VASP) which is concurrently downregulated.

A previous paper reported that GSK3 phosphorylation of Daydreamer, a distantly related orthologue of Lpd, is required to control chemotaxis in *Dictyostelium* ^81^, suggesting that the GSK3-Lpd signalling pathway is evolutionary conserved. From our mass spectrometry screen, we identified multiple GSK3 serine-threonine phosphorylation sites throughout Lpd that fell into two subgroups. First, phosphorylation of Lpd at Thr-1123 overlaps with the Abi1 SH3 domain binding site 2, thus should affect Scar/WAVE binding (Figure 5)^16^. Of the three known Abi binding sites, Abi1 binding site 2 is known to elicit the weakest interaction between Lpd and Abi in the absence of phosphorylation^16^. Indeed, we saw a modest increase in co-immunoprecipitation of the Scar/WAVE complex with Lpd when co-expressed with dominant-active GSK3β (Figure 5). Second, the three serine sites (Ser-1066, Ser-1069, Ser-1071) reside within the Ena/VASP FP4-4 EVH1 domain binding site (Figure 5)^15,80^. GSK3 phosphorylation of these serines should create a negative charge within the core polyproline II helix, potentially reducing interaction with the EVH1 domain. Consistent with this hypothesis, GSK3 activity decreased the binding of Lpd with VASP and Mena (Figure 5). Thus, Lpd may act as an integrator of multiple signals including Rac, PI3-kinase, c-Abl, c-Src, and now also GSK3 to finetune the directed migration of the cell in response to extracellular cues.

Our work also suggests that GSK3 phosphorylates Lpd not only to promote lamellipodia extension but also to prevent focal adhesion maturation. Both GSK3 inhibition and *Lpd* deletion caused a loss of focal complexes underneath the lamellipodium, with the accumulation of mature zyxin- and Mena-positive focal adhesions seen towards the front of cells (Figure 6–7). Similarly, *Lpd* knockout cells have recently been shown to increase mature adhesions at the leading edge of B16-F1 melanoma cells lines^68^. Interestingly, at the whole cell level, the size of zyxin- and Mena-positive focal adhesions was only significantly reduced in *Lpd* heterozygous neural crest cells, compared to wildtype controls and *Lpd* knockout cells (Figure 6–7). This suggests a gene dosage effect of *Lpd* deletion via a secondary currently unknown mechanism.

The subcellular localisation of Ena/VASP proteins is regulated by their interactions of their N-terminal EVH1 domain with FP4 motif-containing proteins^82^, such as Lpd but also RIAM, vinculin and zyxin which recruit Ena/VASP proteins to focal adhesions^83–85^. GSK3-mediated phosphorylation of Lpd may therefore promote the recruitment of Ena/VASP proteins to focal complexes underneath the lamellipodium thereby promoting their conversion to mature focal adhesions. This hypothesis is in agreement with previous observations that Lpd recruits VASP to leading edge clusters, with subsequent budding off of VASP patches which mature into focal adhesions^86^.

Lpd functioning downstream of Rac1 and via Scar/WAVE-Arp2/3 complexes may also contribute to inhibition of focal adhesion maturation since comparable focal adhesion phenotypes to our vinculin immunostaining (Figure 7) have also been seen in Scar/WAVE complex knock-down cells, Arp2/3-depleted melanocytes and *Arp2/3* knockout MEFs^29,69,87^. The peripheral localisation pattern of focal adhesions seen in our *Lpd* knockout neural crest cells was also highly reminiscent of cultured conditional *Rac* knockout neural crest-derived pharyngeal arch cells^88^ and Scar/WAVE complex (*Nckap1)* knockout mouse embryonic fibroblasts^89^. Therefore, the GSK3-Lpd axis together may be required to inhibit the maturation of focal complexes during lamellipodial protrusions, likely by reducing Mena/VASP and increasing Scar/WAVE-Arp2/3 function.

Our study highlights the usefulness of the *ex vivo* experiments to define the GSK3-Lpd axis. However, we do note that the neural crest-specific *Lpd* knockouts did not show any obvious craniofacial phenotypes, in contrast to the neural crest-specific *GSK3* mutants^13^. One reason for the lack of *in vivo* phenotypes could be due to the relative importance of lamellipodia versus filopodia in the *in vivo* neural crest, whereby compensation by filopodia may be sufficient to overcome changes in lamellipodial dynamics^62,63,65,90^. Filopodia can maintain mesenchymal cell persistence by promoting cell-ECM interactions at the cell leading edge and by sensing *in vivo* environmental cues^11,63,65,91^. In agreement, Lpd requires interaction with both actin elongation-associated Ena/VASP proteins and the Scar/WAVE complex during 3D breast cancer invasion^17^.

Moreover, the role of actin regulators during early *in vivo* neural crest development is still contentious. Embryos carrying a neural crest-specific conditional knockout of *Rac1*, *Cdc42* or *FAK* do not show defective *in vivo* neural crest migration, quantified by the number of neural crest cells reaching pharyngeal arch 1, with craniofacial and/or cardiovascular phenotypes only apparent from E11.5-E13.5^88,92,93^. Given that these deletions are in genes transcribing Rho GTPases and enzymes with multiple substrates functioning in actin-based migration, it would therefore follow that Lpd, an actin regulator much further downstream in the actin network, may not show overt phenotypes at early developmental timepoints (E8.5-E9.5).

It is important to note that GSK3 has additional substrates besides Lpd, most notably β-catenin, which is required for neural crest induction and delamination^13,94,95^. The timing of GSK3 pharmacological inhibitor treatment within this study, however, allowed us to bypass GSK3 effects on neural crest delamination and focus on Wnt/β-catenin-independent functions of GSK3 during neural crest migration. GSK3 is also a known cytoskeletal regulator, and can activate proteins such as Rac1 through an unknown mechanism^79,96^ and Rho via phosphorylation of p190ARhoGAP^97^. Focal adhesion kinase (FAK) is also a substrate of GSK3 but how this affects cell migration appears to be more complex: GSK3 phosphorylates FAK at Ser-722 thereby inhibiting FAK kinase activity and consequently reducing cell migration efficiency^78^. However, active GSK3 binds to the phosphodiesterase Prune, which is localised at focal adhesions and is required for FAK and Rac activation, and focal adhesion turnover, which agrees with a positive function in cell migration^79^.

In conclusion, our results suggest that GSK3 serine-threonine phosphorylation of Lpd supports the active migration of neural crest cells, by increasing interactions with the Scar/WAVE complex promoting lamellipodial protrusions, whilst reducing interactions between Lpd and VASP/Mena which drive focal adhesion maturation and filopodia formation. An improved understanding of cytoskeletal regulation in neural crest migration will provide insights into normal development and pathologies such as neurocristopathies and neuroblastoma.

## Supporting information

Dobson Supplementary Movie 1

Dobson Supplementary Movie 2

Dobson Supplementary Movie 3

Dobson Supplementary Movie 4

Dobson Supplementary Movie 5

Dobson Supplementary Movie 6

Dobson Supplementary Movie 7

## ACKNOWLEDGEMENTS

We thank members of the Liu and Krause labs for everyday support, Prof Phillip Gordon-Weeks for insightful discussions, members of CCRB and the NHH Biological Services Unit for ongoing technical assistance.

## AUTHOR CONTRIBUTIONS

Design and management of investigation (MK, KJL), experimental planning and data curation (MK, KJL, LD), collection and analysis of data (LD, WBB, S-YW, ZS, SL), manuscript writing (LD, KJL, MK), review, editing and approval of manuscript (all).

## COMPETING INTERESTS

No competing interests.

## FUNDING

We acknowledge funding from BBSRC BB/R015953/1 (KJL, MK, LD), BB/X512047/1 (KJL, LD), MRC PC21044 (KJL, WBB); MRC-Doctoral Training Programme MRIN013700/1 (LD) and the Double Day Research Fellowship (WBB, KJL).

## STAR METHODS

### KEY RESOURCE TABLE

(See separate document as requested).

### LEAD CONTACT AND MATERIALS AVAILABILITY

Further information and requests for resources and reagents may be directed to and will be fulfilled by the Lead Contacts: Karen J. Liu and Matthias Krause.

### EXPERIMENTAL MODEL AND SUBJECT DETAILS

#### Genetically-Modified Mouse Models

All animal work was approved by King’s College London Ethical Review Process and performed at King’s College London in accordance with UK Home Office Personal License I0DE37907 (LD) and Project Licenses P8D5E2773 (KJL) and PPL9218930 (Julie Keeble). *Tg(Wnt1::*cre*)11Rth* mice previously described in^32^ were crossed with *Lpd* floxed mice (*Raph1^tm1/1Makr^*)^16^. The following Cre-responsive reporters were used, as outlined in figure legends: *LifeAct-EGFP (Tg(CAG-EGFP)#Rows),* which encodes a fluorescent reporter of filamentous actin dynamics^31^, and *Rosa26R^mtmg^* (*GT(Rosa)R26Sor^Tm4(ACTB-tdTomato-EGFP)Luo^),* whereby a membrane tdTomato-polyA is flanked by loxP sites followed by a membrane-EGFP in the Rosa26 locus, which encodes plasma membrane dynamics^30^. All mouse lines were bred on an outcrossed CD1 background. Mice were genotyped as described in original publications. Gestational ages were determined based on the observation of vaginal plugs, which was considered E0.5. Embryos were further staged by determining somite stage after dissection. For each experiment, litter-matched controls were used unless otherwise noted.

#### Embryonic Dissections

At days corresponding to embryonic day E8.5 or E9.5, the mother was sacrificed, and her uterus dissected out and immediately placed into ice cold 1x PBS. In a 10 cm^2^ dish, the mesometrium of the uterus was cut, and the muscle layer removed to separate out individual decidua. The decidual tissue was peeled back and extra-embryonic tissues removed. E8.5 and E9.5 embryos were transferred into a 24-well tissue culture plate for fixation (see Method Details).

#### Live E8.5 Embryo Culture

*Wnt1*::cre; *Rosa26R^mtmg^* E8.5 embryos (6-10 somite stage) were dissected out from the decidua of the mother’s uterus, and the extra-embryonic membranes removed. The embryos were transferred into a 35 mm coverslip glass-bottomed dish (Ibidi) and maintained in culture media (Dulbecco’s modified Eagle’s medium (DMEM) High Glucose (phenol red-free) (Sigma), 50% rat serum (Envigo)) and incubated at 37°C and 5% CO_2_. *Wnt1*::cre-positive embryos expressing membrane-EGFP (mGFP) in the neural crest domains, as determined using an epifluorescent lamp attachment to the dissection stereoscope, were taken forward for live imaging. Embryos were maintained in culture for a maximum of 5-6 hours prior to fixation.

#### Primary Neural Crest Explant Cultures

The full method of this protocol can be found in^33^. Briefly, embryonic day 8.5 (E8.5) mouse embryos were dissected out from the decidua of the mother’s uterus, and extra-embryonic membranes removed. The head fold was removed from the body of the embryo, at an anteroposterior position just anterior to the heart (see Figure 2A). The underlying mesoderm beneath the neural plate border (NPB) was scraped away and the cleaned NPB divided down the anteroposterior axis so that each side of the neural plate border could be plated individually. Using a glass Pasteur pipette, the NPB was transferred into coverslip-bottomed 24-well plates (Ibidi). Each well was pre-coated with 1 µg/ml fibronectin (Sigma) and the explants were cultured in neural crest media (Dulbecco’s modified Eagle’s medium (DMEM)-high glucose (Sigma), 15% embryonic stem cell-grade foetal bovine serum (Sigma), 0.1 mM minimum essential medium nonessential amino acids (Gibco), 1 mM sodium pyruvate (Sigma), 55 µM β-Mercaptoethanol (Gibco), 100 units/mL penicillin, 100 units/mL streptomycin and 2 mM L-Glutamine, conditioned by growth-inhibited STO feeder cells (ATCC) and supplemented with 25 ng/μl basic-FGF (R&D Systems) and 1000 U LIF (ESGRO by Millipore) and incubated overnight at 37°C and 5% CO_2_. The outgrowth of the pre-migratory and migratory neural crest cell populations is visible by 24 hours following dissection.

#### Human Cell Lines

HEK293FT cells (Thermo Fisher) were cultured in high-glucose Dulbecco’s modified Eagle’s medium (DMEM) (Sigma) supplemented with 10% of foetal bovine serum (FBS, Gibco), 2 or 4 mM L-glutamine, 1 unit/ml penicillin and 100 μg/ml streptomycin. Cells were maintained in 75 cm^2^ or 175 cm^2^ tissue culture flasks (Greiner) and incubated at 37°C and 10% CO_2_.

### METHODS DETAILS

#### Molecular Biology and Transfections

The following materials were used: pBSII KS-(Agilent Technologies), pEGFP-N1 (Clontech), pDONR221-WAVE-2 (German Resource Centre for Genome Research). Sra1 (CYFIP1; HsCD00042136), Nap1 (NCKAP1; HsCD00045562) and HSPC300 (C3orf10; HsCD00045008) (DNASU repository) in pENTR233 or pDONR221. hsAbi1d (BC024254; Geneservice) full-length was cloned into pENTR11 (Invitrogen)^16^. pMyc-VASP, pMyc-Mena and pMyc-EVL were cloned by PCR amplification of murine VASP/Mena/EVL cDNA from existing plasmids using primers (listed in: Key Resource Table) into pENTR3C (Invitrogen). Scar/WAVE and Ena/VASP cDNAs were transferred to pRK5-myc-DEST (kind gift of Jean-Paul Borg, Marseille Cancer Research Centre, France) by Gateway® cloning for CMV-driven expression in mammalian cells. Human Lpd (AY494951) was amplified and cloned into pENTR3C (Invitrogen) and transferred to the pCAG-DEST-EGFP mammalian expression vector using Gateway® recombination^17^. HA GSK3 beta S9A pcDNA3 was a gift from Jim Woodgett (Addgene plasmid # 14754).

HEK293FT cells (Thermo Fisher) were transiently transfected using Lipofectamine 2000 (Invitrogen) according to manufacturer’s instructions, branched Polyethylenimine (PEI) (Sigma) or calcium phosphate. Lipofectamine 2000 or branched PEI (Sigma) were used for small-scale transfections: 1 x10^6^ HEK293FT cells were transfected with 4 µg of the following plasmids: pCAG-Lpd-EGFP, pEGFP-N1, pBSII KS-, pCDNA-hsS9A-GSK3β-HA, pMyc-VASP, pMyc-Mena, pMyc-EVL, pMyc-Abi-1D, pMyc-WAVE2, pMyc-Sra-1, pMyc-Nap1 and pMyc-HSPC300. For PEI-based transfection, 4 μg of DNA and 8 μl of PEI was separately diluted in 100 μl OptiMEM. The solutions were flick mixed and incubated at room-temperature for 5 min. The OptiMEM-PEI mix was added to the Opti-MEM DNA mix and incubated at room temperature for 20 min. The transfection mix was added dropwise to cells and incubated at 37°C and 10% CO_2_ for 24h prior to cell lysis. Calcium phosphate transfection: 12 x10^6^ cells were transfected with 50 µg of the following plasmids: pCAG-Lpd-EGFP, pEGFP-N1, pBSII KS-, pCDNA-hsS9A-GSK3β-HA. Briefly, 1.25 M CaCl_2_ was added to the DNA solution. 2x HBS solution (50mM HEPES pH 7.05, 10 mM KCl, 12 mM Dextrose, 280 mM NaCl, 1.5 mM Na_2_HPO_4_) was added dropwise to the CaCl_2_-DNA mix with bubbling. The transfection mix was added dropwise to cells, before being replaced with fresh media 3.5-4 hours later. HEK293FT cells were harvested 48 hours after transfection.

#### Pharmacological Inhibitor Treatments

The competitive pharmacological inhibitors of GSK3α/β, 6-bromoindirubin-3’-oxime (BIO, Sigma) and CHIR99021 (Tocris Bioscience), were re-suspended in dimethyl sulphoxide (DMSO, Sigma) at a stock concentration of 14 mM and 10 mM, respectively. The inhibitors were further diluted in standard cell media to final working concentration. HEK293FT cells were treated with 1 µM BIO for 24 hours prior to cell lysis. Neural crest explant cultures were treated with DMSO vehicle control, 0.5 µM BIO or 1 µM CHIR99021 for 2 hours or 24 hours prior to fixation. For drug rescue experiments, neural crest explants were treated with BIO for 2 hours, washed twice with 1x PBS and fresh neural crest media added for the set time stated in the figure legends prior to fixation.

#### Tissue Fixation

Wholemount E8.5 and E9.5 embryos were fixed in ice-cold 4% paraformaldehyde (PFA) in PBS overnight with gentle rocking. The following day, the PFA was removed and 3x 15 min 1x PBS washes completed. Neural crest explant cultures were fixed 48 hours after dissection with 4% paraformaldehyde (PFA)-PHEM (60 mM PIPES, 25 mM HEPES, 10 mM EGTA, 2 mM MgCl_2_, 0.12 M sucrose) for 10 minutes at room temperature.

#### Tissue Processing for Cryosectioning

Following their dissection and fixation, E9.5 embryos were transferred into graded sucrose solutions for cryoprotection. Firstly, samples were incubated in 30% sucrose-PBS solution at 4°C overnight, followed by a subsequent overnight 4°C incubation in 30% sucrose-OCT (Optimal Cutting Temperature) solution (CellPath). Once equilibrated, the E9.5 embryos were anteriorly embedded into OCT blocks and snap frozen using dry ice and 100% ethanol. An OTF-5000 Cryostat (Bright), set to −16°C specimen temperature and −24°C chamber temperature, was used to section the cryo-blocks (14 μm thickness). Cryosections were sequentially mounted over two SuperFrost-Plus^®^ glass slides (Thermo Fisher) and stored at −80°C.

#### Immunofluorescence

Wholemount embryo immunofluorescence: E8.5 and E9.5 embryos were permeabilised in 0.5% Triton-X-100-PBS at room temperature. Embryos were blocked in 10% goat serum-0.1% Tween 20-PBS at 4°C for 24-48 hours, prior to incubation with primary antibody diluted in blocking buffer at 4°C for 24 hours. Samples were washed with blocking buffer at room temperature before being incubated with Alexa488-conjugated secondary antibodies and Hoechst at 4°C for 24 hours. The samples were placed into Citifluor (50% glycerol anti-fade mounting media) to clear at 4°C for 2 days and 5 days, for E8.5 and E9.5, respectively.

Cryosection immunofluorescence: slides were washed with 0.1% Triton-X-100-PBS in a coplin jar to remove any remaining OCT, and the perimeter of the slide outlined using an ImmEdge™ hydrophobic barrier pen (Vector Labs). The sections were blocked in 10% normal goat serum-1% BSA-0.1% Triton-X-100-PBS for 1 hour at room temperature, before being incubated with a chicken anti-EGFP primary antibody diluted to 1:500 in blocking buffer at room temperature for 1 hour. The slides were subsequently washed and incubated with Alexa488- or Alexa568-conjugated secondary antibodies diluted 1:400 in blocking buffer at room temperature for 1 hour. The slides were washed, and a coverslip mounted over the samples using Fluoroshield mounting medium with DAPI (Abcam).

Neural crest explants were permeabilised at room temperature for 2 min with 0.1% Triton-X-100 in cytoskeletal-(c)TBS buffer (200 mM Tris-HCl, 1.54 M NaCl, 20 mM EGTA, 20 mM MgCl_2_ x 6H_2_0 pH 7.5), before being blocked overnight at 4°C (10% normal goat serum-10% BSA-cTBS buffer). Alternatively, the explants were permeabilised with 0.05% saponin as part of the blocking buffer and incubated overnight at 4°C. The samples were incubated with primary antibodies diluted in 1% BSA-cTBS for 1 hour at room temperature, followed by 3x cTBS washes and incubation with secondary antibodies diluted in 1% BSA-cTBS for 1 hour at room temperature. Nuclei were stained with 1:1000 dilution of Hoechst 33342 (20 mg/ml stock concentration) as part of the secondary antibody mix. For coverslip immunofluorescence samples, coverslips were mounted onto SUPERFROST^®^ microscope slides (Thermo Fisher) with Fluoroshield Mounting Medium with DAPI (Abcam).

#### VASP Monoclonal Antibody Production

His-tagged full length murine VASP was produced in insect cells using the “Bac-to-Bac” baculovirus expression system according to the protocols supplied by the manufacturer (GIBCO-BRL) and purified on cobalt beads (BD Talon resins, BD Biosciences Clontech). These recombinant proteins were used to produce monoclonal antibodies in VASP knockout mice as described^98^. Hybridoma supernatants were screened by ELISA on recombinant, purified VASP and on western blots of extracts of Swiss 3T3 fibroblasts. One hybridoma was chosen and subcloned twice. This monoclonal antibody was designated B296B5H12 and recognises VASP of murine origin.

#### Pulldowns and Western Blotting

HEK293FT cells were lysed on ice for 15 min using glutathione S-transferase (GST) buffer (50 mM Tris-HCl pH 7.4, 200 mM NaCl, 1% NP-40, 2 mM MgCl_2_, 10% glycerol) supplemented with NaF (10 mM final concentration), Na_3_VO_4_ (1 mM final concentration) and a complete protease inhibitor mini tablet (Roche). Samples were centrifuged at 4°C for 10 min. Protein concentration was determined using a Pierce BCA Assay (Thermo Fisher). 400 μg protein was incubated on pre-blocked (1% BSA-GST buffer) GFP-Trap® (Chromatek) or GFP-Selector beads (NanoTag) at 4°C for 2 hours. Pulldown samples were washed and re-suspended in 2x sample buffer (100 mM Tris-HCl (pH 6.8), 4% SDS, 12% glycerol, 4 mM DTT, 0.02% bromophenol blue), and run alongside 20 μg lysate and peqGOLD Protein Marker V (VWR International) on 10% SDS-PAGE gels (stacking gel: 5% bis-acrylamide (30%), 125 mM Tris-HCl (pH 6.8), 0.1% SDS, 0.1% APS, 0.05% TEMED); separating gel: 10% bis-acrylamide (30%), 400 mM Tris-HCl (pH 8.8), 0.1% SDS, 0.1% APS, 0.05% TEMED). Western blot transfer onto Immobilon PVDF membranes (EMD Millipore) was then completed: 100 V, 350 mA, 17 W for 1.5 hours. Membranes were blocked at 4°C overnight in 5% BSA-TBS-T (20 mM Tris-Base, 154 mM NaCl, 0.1% Tween-20, pH 7.6) and subsequently incubated at room temperature for 1h with primary antibodies, followed by washes and a further 1 hour with HRP-conjugated secondary antibodies (Cell Signalling Technology). Blots were washed and developed with the ECL Western Blotting Detection Kit (Bio-Rad Laboratories) and imaged using a Bio-Rad Imager and ImageLab software. Western blots were quantified using the pixel densitometry tool (FIJI/ImageJ). The profile plot for each western blot lane was generated and the area underneath the graph, representing the relative band density, was calculated and normalised to the pixel density of the Lpd-EGFP pulldown lane.

#### Phosphorylation Analysis by Tandem Mass Spectrometry

HEK293FT cells were transiently transfected with the following plasmids: Lpd (45 µg pCAG-Lpd-EGFP, 5 µg pBSII KS-), Lpd + BIO (45 µg pCAG-Lpd-EGFP, 5 µg pBSII KS-), Lpd + GSK3b (45 µg pCAG-Lpd-EGFP, 5 µg pCDNA3-hsS9A-GSK3b-HA) prior to cell lysis. Protein concentration was quantified and the samples incubated with pre-blocked (1% BSA-GST buffer) GFP-Trap® beads for 2 hours at 4°C for pulldown. Pulldown samples were re-suspended in 2x sample buffer (100 mM Tris-HCl (pH 6.8), 4% SDS, 12% glycerol, 10 mM DTT, 0.05% bromophenol blue), and run on 6% Novex WedgeWell™ Tris-Glycine mini protein gels at 225 V for 45 min before Colloidal Coomassie staining (Severn Biotech). Following de-staining (HPLC-grade water), the protein band corresponding to Lpd-EGFP (approximately 230 kDa) was excised and in gel digestions completed using a tri-enzyme mix (trypsin-chymotrypsin-AspN). Collision-induced dissociation (CID) / electron transfer dissociation (CID/ETD) tandem mass spectrometry was then performed. Chromatographic separation was completed using a U3000 UHPLC NanoLC system (Thermo Fisher). Peptides were resolved by reversed phase chromatography on a 75 μm C18 Pepmap column (50 cm length) using a three-step linear gradient of 80% acetonitrile in 0.1% formic acid. The gradient was delivered to elute the peptides at a flow rate of 250 nl/min over 60 min starting at 5% B (0-5 minutes) and increasing solvent to 40% B (5-40 minutes) prior to a wash step at 99% B (40-45 minutes) followed by an equilibration step at 5% B (45-60 minutes). The eluate was ionised by electrospray ionisation using an Orbitrap Fusion Lumos (Thermo Fisher) operating under Xcalibur v4.3. The instrument was first programmed to acquire using an Orbitrap-Ion Trap method by defining a 3 second cycle time between a full MS scan and MS/MS fragmentation by collision induced dissociation. Orbitrap spectra (FTMS1) were collected at a resolution of 120,000 over a scan range of m/z 375-1600 with an automatic gain control (AGC) setting of 4.0e5 (100%) with a maximum injection time of 35 ms. Monoisotopic precursor ions were filtered using charge state (+2 to +7) with an intensity threshold set between 5.0e3 to 1.0e20 and a dynamic exclusion window of 35 seconds ± 10 ppm. MS2 precursor ions were isolated in the quadrupole set to a mass width filter of 1.6 m/z. Ion trap fragmentation spectra (ITMS2) were collected with an AGC target setting of 1.0e4 (100%) with a maximum injection time of 35 ms with CID collision energy set at 35%. Neutral loss scans were performed to trigger fragmentation in the presence of phosphorylation with simultaneous triggering of ETD fragmentation scans. Data processing and analysis was completed in Proteome Discoverer v2.5 with the .msf files uploaded in to Scaffold 5 for manual interpretation of MS/MS fragmentation spectra and site localisation.

#### Live embryo imaging: Protrusion dynamics

Live imaging of *Wnt1*::cre; *Rosa26R^mtmg^* embryos was performed on a Nikon A1R inverted confocal microscope with an environmental chamber set to 37°C and 5% CO_2_. The E8.5 embryos were maintained and positioned laterally in phenol red-free culture medium in 35 mm coverslip glass-bottomed dishes (Ibidi) which were mounted onto the microscope. Embryos were located and oriented using the 488 nm emission laser at 10x magnification. Live imaging focused on those mGFP-positive neural crest cells delaminating and early emigrating away from the neural plate border, destined for pharyngeal arch 1. Live imaging was completed at 40x magnification at a z-depth of 40 μm, over 30 min (1 frame/ 45 sec) or at a z-depth of 24.5 μm, over 10 min (1 frame/ 20 sec).

#### Live cell imaging: Neural crest migration and protrusion dynamics

Live imaging of neural crest migration was completed 24 hours after dissection on a widefield IX 81 microscope (Olympus), with a Solent Scientific incubation chamber (37°C; 5% CO_2_), filter wheels (Sutter), an ASI X-Y stage, Cascade II 512B camera (Photometrics), and 4x UPlanFL, 10x UPlanFL, 60x Plan-Apochromat NA1.45, or 100x UPlan-Apochromat S NA 1.4 objective lenses, controlled by MetaMorph software. Lineage-labelled neural crest cells were located using the 488 nm filter from a Xenon white light source at 10x magnification, and the explants oriented so that the neural plate border was just outside the frame of imaging. Phase-contrast live imaging was then completed at 10x magnification, over 18 hours (1 frame/ 5 min). A minimum of two regions of interest (ROIs) were imaged per explant and multi-well imaging was performed using an ASI x-y stage to capture the same time intervals across genotypes or between drug conditions. Following live imaging, the time-lapse movies were exported from the MetaMorph software and saved as TIFF stacked (stk) files for downstream analysis.

Live imaging of protrusion dynamics of *Wnt1*::cre; *LifeAct-EGFP* migratory neural crest cells were completed 36 hours after dissection on an IX 81 widefield epifluorescence microscope (Olympus), with a Solent Scientific incubation chamber (37°C; 5% CO_2_), controlled by MetaMorph software. Lineage-labelled neural crest cells were located using the 488 nm filter from a Xenon white light source at 10x magnification. Imaging focused on LifeAct-EGFP-expressing migratory neural crest cells at the explant edge. Live imaging was completed at 60x magnification, over 10 min (1 frame/ 5 seconds). 3 cells were imaged per explant, and three biological repeats performed.

#### Imaging: Fixed Immunofluorescence

Wholemount E8.5 and E9.5 embryos were imaged on an inverted Nikon A1R confocal microscope. Low magnification (4x) images were acquired as z-stacks with 40 μm slice interval (800 μm). High magnification (20x) images were acquired as z-stacks with 2μm slice interval (80-100 μm). Cryosection slides were imaged on a Leica TCS SP5 DM16000 confocal microscope at 20x and 63x magnification (with 1x or 2x optical zoom). Z-stacks were taken at 1 μm intervals through 14 μm tissue sections.

Neural crest explants were imaged on a widefield epifluorescence IX81 Olympus microscope (Olympus), with a Solent Scientific incubation chamber (37°C; 5% CO_2_), filter wheels (Sutter), an ASI X-Y stage, Cascade II 512B camera (Photometrics), and 4x UPlanFL, 10x UPlanFL, 60x Plan-Apochromat NA1.45, or 100x UPlan-Apochromat S NA 1.4 objective lenses, controlled by MetaMorph software. Lineage-labelled neural crest cells were located using the 488 nm filter from a Xenon white light source at 10x magnification. Imaging was completed at 60x magnification, with equal exposure times used for all cells and conditions imaged for a given experiment.

#### Quantification of migration speed and persistence

Migratory cranial neural crest cells were manually tracked through the course of time-lapse by following cell nuclear position using the Manual Tracking plugin (ImageJ/Fiji). 10 lineage-labelled neural crest cells were tracked per explant, specifically those within the 2 most outward rows of mesenchymal cells at the explant edge. Cell tracks were stopped prematurely if cells underwent division or if they left the frame of imaging. Cell tracking generated XY coordinates over time which were exported into Microsoft Excel. XY coordinates were subsequently converted into matrix format using the “Convert ImageJ files into .CEL format” script (Mathematica) for import into the Chemotaxis Analysis Notebook v1.5β (Mathematica) to analyse mean track speed (MTS) (G. Dunn, King’s College London). MTS is defined as the distance travelled by the neural crest cells over a set time ratio (TR), averaged across the entire track length^16,99^. MTS is calculated as an average displacement (*d_n_*) over dt, where n denotes which track interval (n = 1 indicates 5 min) and dt is the usable time interval (dt = 5 min)^16,99^.

Mean squared displacement (MSD) is a speed-and persistence-dependent measure of the area explored by cells over a set time^34^. Direction autocorrelation is a speed-independent measure of cell directionality, through the calculation of how the angle of displacement vectors correlate with themselves^34^. Quantification of MSD and direction autocorrelation were calculated using the excel macros provided in^34^, according to the protocol provided.

#### Quantification of Cell Morphology and Actin-based Protrusions

Fixed migratory neural crest cells from E8.5 *Wnt1*::cre; *LifeAct-EGFP* explant cultures (with or without *Lpd knockout* or GSK3 inhibition) were manually outlined and masked by a blinded examiner using the LifeAct-EGFP fluorescence to define the cell edge (Freehand Selection tool, ImageJ/FIJI). The cell area and circularity were calculated using the Analyse | Measure function (ImageJ/FIJI). Manual filopodia counts were completed using the LifeAct-EGFP fluorescence, the number of which were first normalised against the individual neural crest cell area and then multiplied by either a factor of 100 (*in vivo* datasets) or 5000 (*ex vivo* datasets) to generate filopodia number per 100 or 5000 µm^2^.

#### Quantification of leading edge actin-associated protein localisation

Fixed migratory neural crest cells stained for Abi1, Lpd and Mena (568 nm filter) were merged with their respective actin reporter, LifeAct-EGFP, image (488 nm filter) using ImageJ/FIJI. A blinded examiner classified neural crest cells as having either a positive or negative actin-associated protein stain localised (LifeAct-EGFP) at the lamellipodia edge. A positive stain was classed as continuous staining of the actin-associated protein at the lamellipodium. A compromised stain was classified as punctate protein localisation or evidence for membrane ruffling. Filopodia or an absence of leading edge staining were classified as negative. Data was presented as the percentage of cells with positive, compromised or negative staining for the actin-associated protein at the leading edge ± standard error of the mean (SEM).

#### Quantification of focal adhesion protein localisation

Fixed migratory neural crest cells stained for Mena, vinculin or zyxin were manually masked using the Freehand Selection Tool | Fit Spline in ImageJ/FIJI by a blinded examiner. For cell third measurements (front, middle, back), neural crest cells were divided according to 1/3 cell area down the major axis of the cell, relative to the overall direction of migration. Manual cell 1/3 masks were generated as above. The masked TIFF images were imported into the Focal Adhesion Analysis Server (FAAS) (see Key Resource Table) and the detection threshold (DT, standard deviation) and the minimum pixel size (MPS, µm) set: Mena: DT 2, MPS 3; vinculin: DT 2.5, MPS 5; zyxin: DT 2.5, MPS 5). Optimisation of the detection threshold and minimum pixel size was completed prior to analysis using a training data set. Output measurements for number, area, length, and width of focal adhesions were exported into Microsoft Excel and the mean of each parameter averaged per cell. Filopodia number was normalised to a standardised area of 100 µm^2^.

### QUANTIFICATION AND STATISTICAL ANALYSIS

Statistical analysis was performed in Prism v8 or v9 (GraphPad) using a Student’s t-test, one-way ANOVA with Tukey’s multiple comparison’s test, or chi-squared test (see figure legends). P values < 0.05 were considered significant.

### DATA AND CODE AVAILABILITY

The proteome data will be deposited on an appropriate repository.

## SUPPLEMENTAL INFORMATION

(See separate document as requested).

## Notes

### Competing Interest Statement

The authors have declared no competing interest.

