## Supplemental Figures and Legends for "GSK3 and Lamellipodin balance lamellipodial protrusions and focal adhesion maturation in mouse neural crest migration"

Supplementary Figures

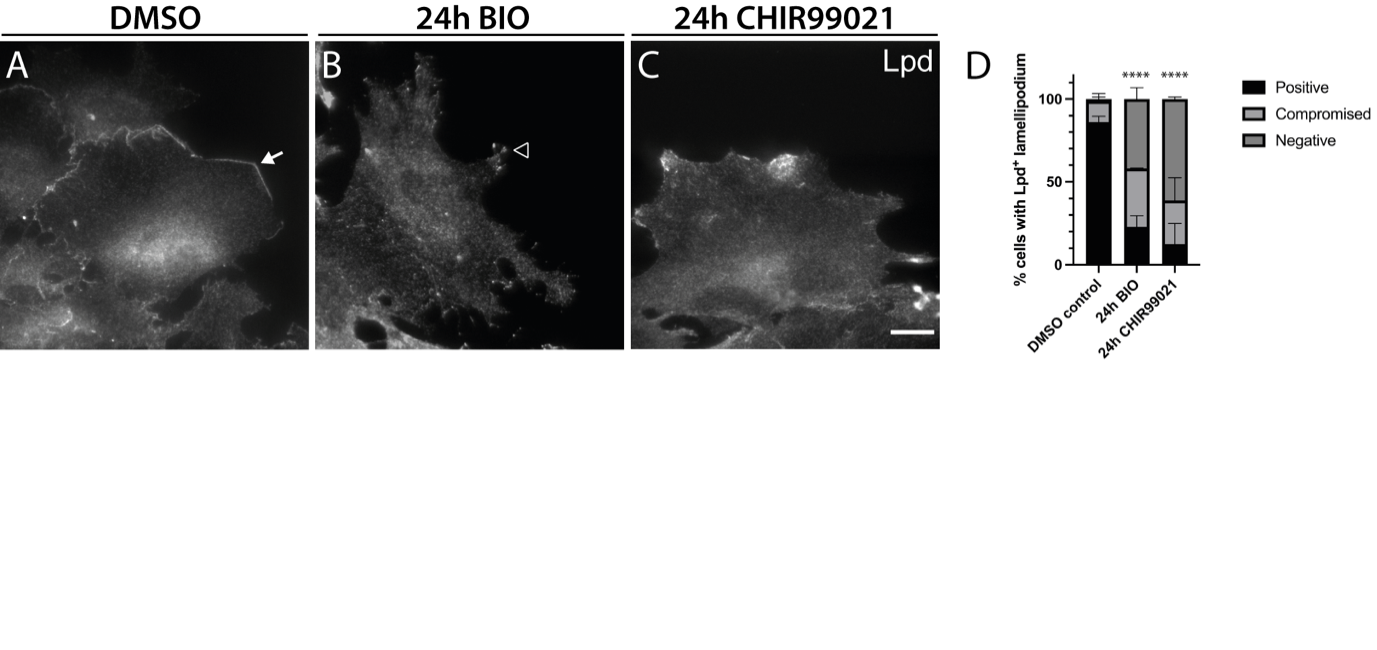

**Supplementary Figure 1.** **Long-term GSK3 inhibition mislocalises** **Lpd away from the leading edge of mouse cranial neural crest cells. (A-C)** Lpd immunostaining of fixed migratory cranial neural crest cells, cultured from CD1 WT E8.5 embryos, treated for 24 hours with **(A)** DMSO control, **(B)** 0.5 µM BIO or **(C)** 1 µM CHIR99021. Scale bar 20 µm. Arrow: continuous Lpd immunostaining, arrowhead: Lpd localisation at filopodial tips. **(D)** Quantification of the percentage migratory neural crest cells with continuous Lpd-labelled lamellipodium (positive), a discontinuous Lpd-labelled lamellipodium (compromised) or absence of Lpd-labelled lamellipodium after 24 hour treatment with DMSO or the GSK3 inhibitors, BIO and CHIR99021. Data presented as mean ± SEM, and analysed over at least 3 independent experiments. **** p < 0.0001, chi-squared test. DMSO control: n= 78 cells, 24h BIO: n= 57 cells, 24h CHIR99021: n= 31 cells.

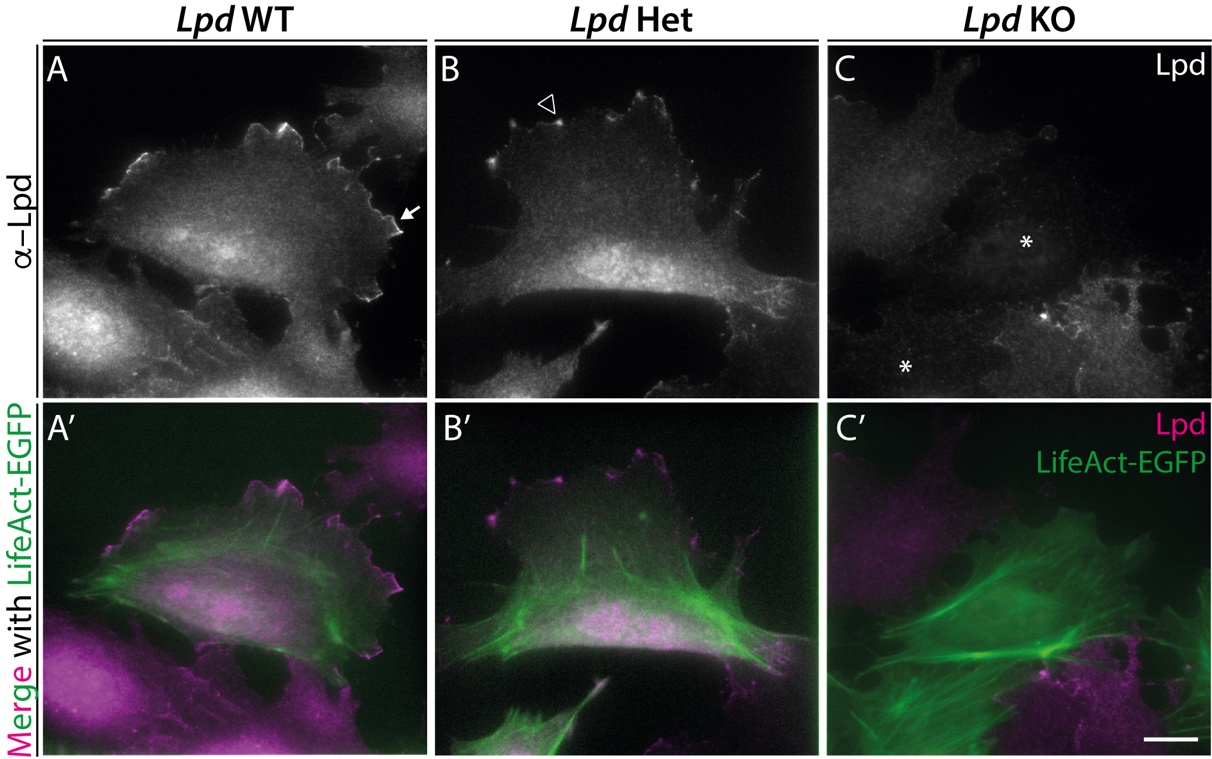

**Supplementary Figure 2.** **Lpd protein is lost from *Lpd* knockout cranial neural crest cells two days after *Wnt1*::cre deletion. (A-C)** Lpd immunostaining of fixed migratory cranial neural crest cells, cultured from E8.5 *Wnt1*::cre; *Lpd*^+/+^ **(A)**, *Wnt1*::cre; *Lpd*^+/fl^ **(B)**, and *Wnt1*::cre; *Lpd*^fl/fl^ **(C)** embryos. **(A’-C’)** The actin filaments of *Wnt1*::cre-expressing neural crest cells are genetically labelled with LifeAct-EGFP (green) as seen in the merged images with Lpd (magenta), which was also used to identify neural crest cells within the explant cultures. Arrow: continuous Lpd immunostaining, arrowhead: punctate Lpd localisation, *: loss of Lpd. Scale bar: 20 μm.

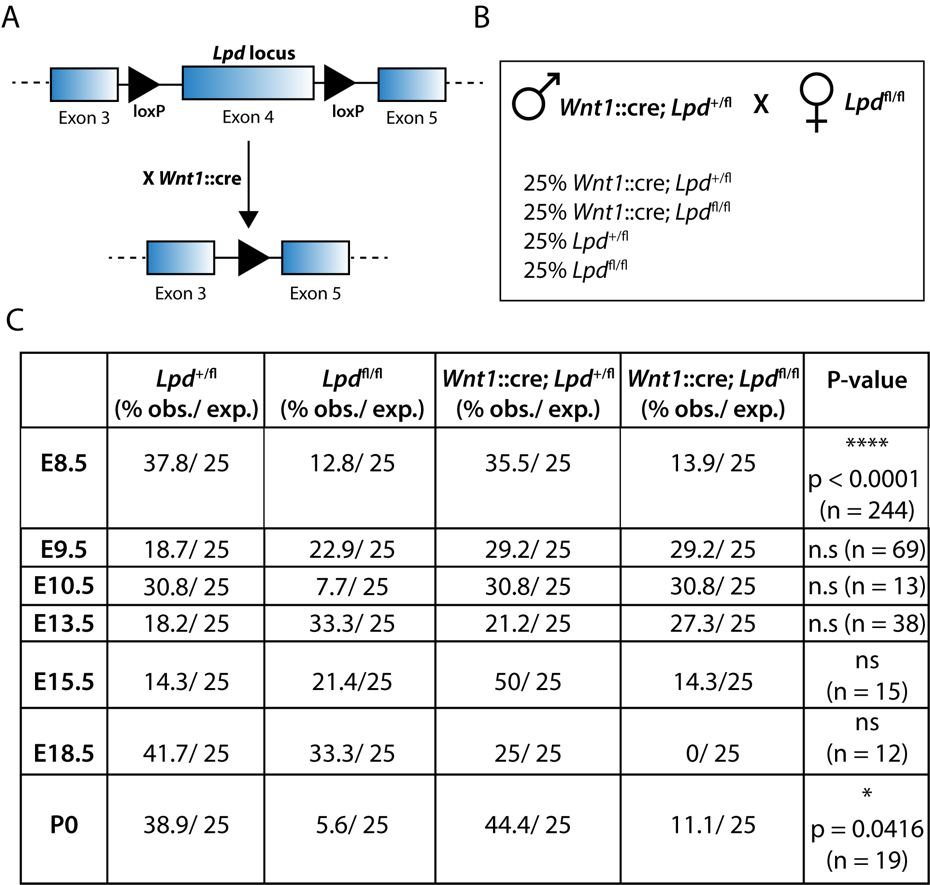

**Supplementary Figure 3. Distribution of genotypes seen upon the neural crest-specific deletion of *Lpd*. (A)** Schematic representation of the *Lpd* conditional knockout mouse model used in this study, which excises exon 4 of the *Lpd* (*RAPH1*) gene to cause premature truncation and loss of function (Law et al., 2013). **(B)** Parental cross used to obtain conditional *Lpd* heterozygous and *Lpd* homozygous knockout embryos. **(C)** Table of the observed and expected mendelian ratios of embryonic day (E) 8.5, E9.5, E10.5, E13.5, E15.5, E18.5 embryos and P0 pups obtained from the parental cross in **(B)**. The observed percentage of *Wnt1*::cre; *Lpd*^fl/fl^ embryos was significantly reduced compared to expected ratios at E8.5 as well as at the early post-natal time-point, P0. The intervening timepoints might not show statistical significance due to the low numbers analysed. A chi-squared test was used to quantify the differences between the observed versus expected genotypes over developmental stages. **** p < 0.0001, * p < 0.05, ns non-significant.

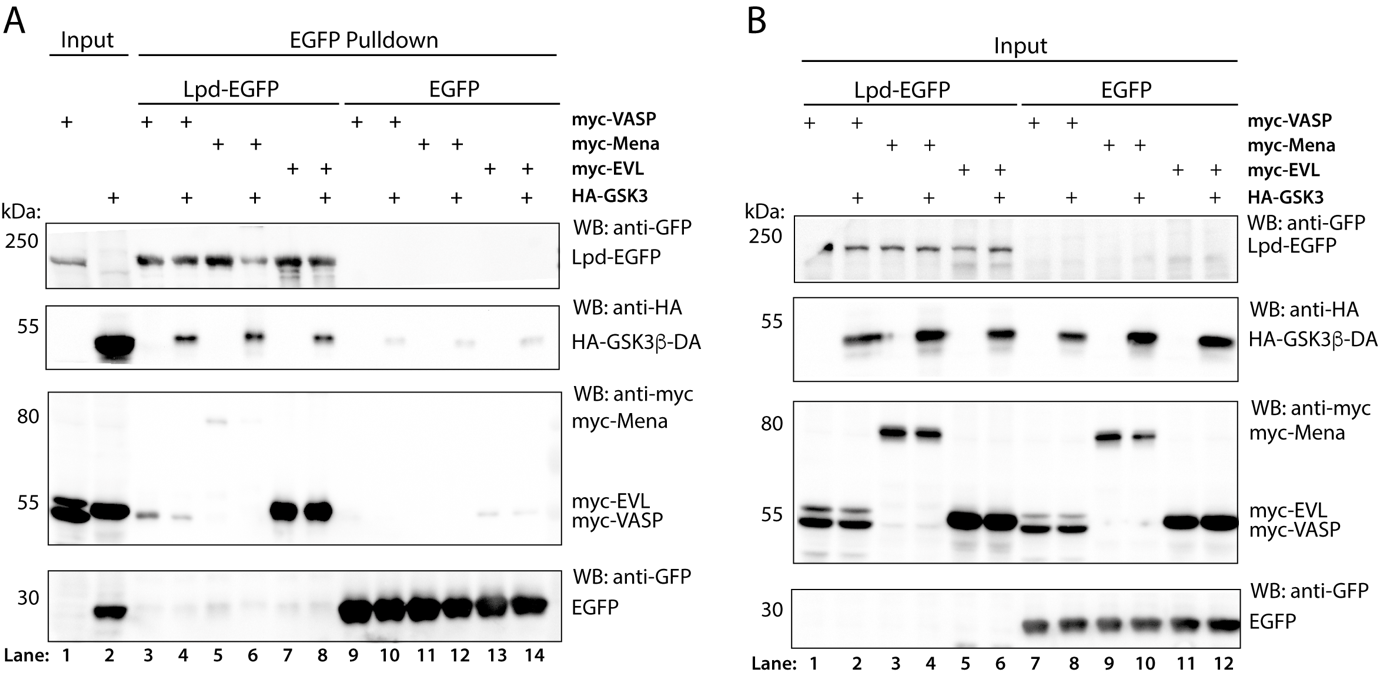

**Supplementary Figure 4. GSK3 kinase activity reduces the interaction of Lpd with the Ena/VASP proteins, VASP and Mena.** HA-tagged, dominant-active (DA) GSK3β (DA-GSK3β-HA) co-immunoprecipitation with Lpd-EGFP and myc-tagged En/VASP proteins (myc-VASP, myc-Mena, myc-EVL) in HEK293FT cells. **(A)** EGFP-Trap pulldowns were performed from cell lysates followed by western blotting. This panel **(A)** is a repeat of (**Figure 5E**), shown here for completeness. **(B)** Lpd-EGFP, DA-GSK3β-HA, myc-VASP, myc-Mena, myc-EVL and EGFP protein overexpression was confirmed by running protein lysate samples on a western blot, parallel to EGFP pulldown samples. Blots were probed for anti-myc, anti-HA and anti-EGFP, and are representative of 3 independent experiments.

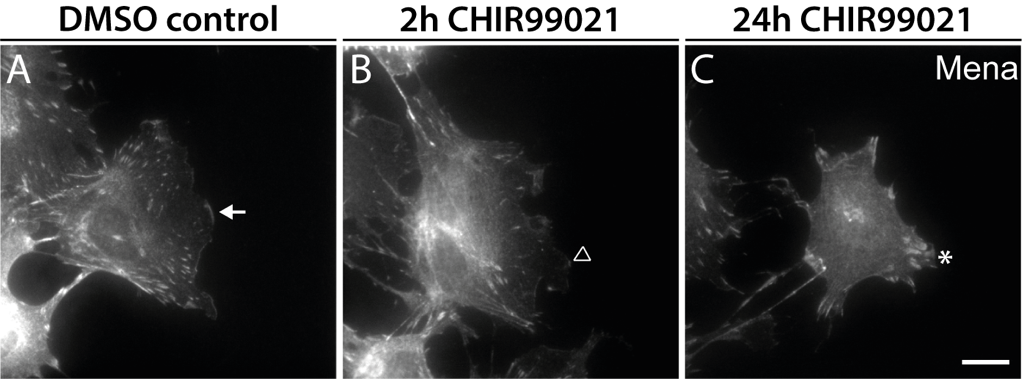

**Supplementary Figure 5. GSK3 inhibition (CHIR99021) reduces Mena leading edge localisation and increases Mena-positive focal adhesions in neural crest cells. (A-C)** Mena immunostaining of fixed migratory cranial neural crest cells, cultured from E8.5 WT embryos treated with **(A)** DMSO vehicle control or CHIR99021 for **(B)** 2 hours or **(C)** 24 hours prior to fixation. **(A)** In DMSO control cells, Mena is localised to the edge of lamellipodia (arrow). **(B)** Short-term CHIR99021 treatment ablates the lamellipodia, with Mena localisation to filopodia and focal adhesions (open arrowhead), which is exacerbated with 24 hour treatment (**C**, *). For quantification, see (**Figure 6P, S-T**). Scale bar: 20 μm.

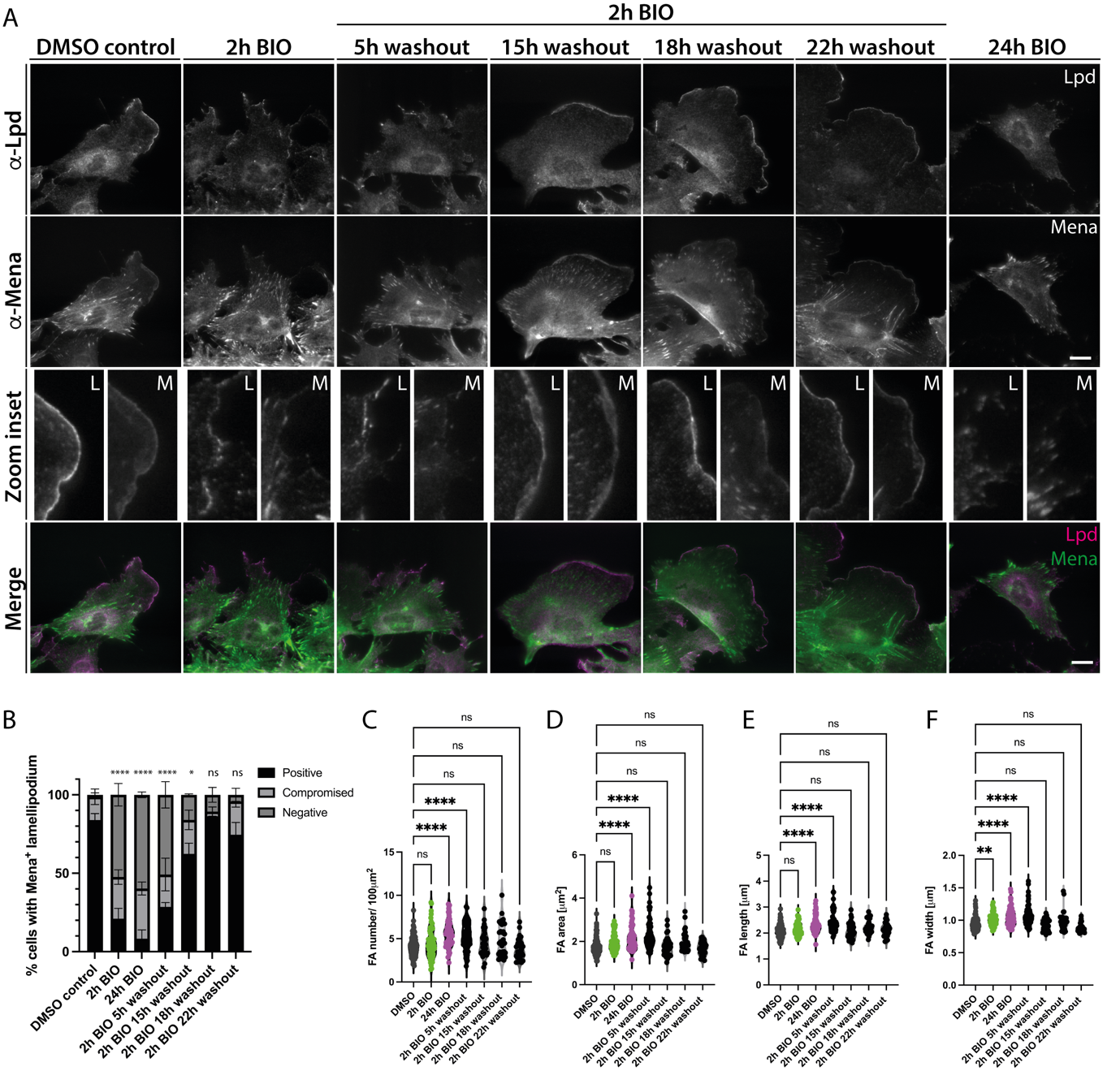

**Supplementary Figure 6. GSK3 is required for the rapid and reversible regulation of Lpd-Mena localisation in primary neural crest cells. (A)** Lpd and Mena co-immunostaining of fixed migratory cranial neural crest cells, cultured from E8.5 WT embryos, treated with DMSO control or GSK3 inhibitor, BIO, for 2 hours. Following 2h BIO treatment, explants were either fixed, or subjected to a drug wash-out whereby fresh media was applied to the explants which were allowed to recover for 5h, 15h, 18h or 22h prior to fixation. A 24h BIO treatment was also included. **(B)** Quantification of the percentage neural crest cells with a Mena-positive lamellipodium, plotted as mean ± SEM, **** p < 0.0001, * p < 0.05, ns: non-significant, chi-squared test. **(C-F)** Quantification of Mena-positive **(C)** focal adhesion number/ 100μm^2^, **(D)** focal adhesion area, **(E)** focal adhesion length and **(F)** focal adhesion width of cranial neural crest cells across the drug treatment conditions. ns: non-significant, ** p<0.01, *** p<0.001, **** p<0.0001, one-way ANOVA with Tukey’s multiple comparisons test. DMSO control: N=99 cells, 2h BIO: N=79 cells, 24h BIO: N=61 cells, 2h BIO 5h washout: N=57 cells, 2h BIO 15h washout: N=31 cells, 2h BIO 18h washout: N=30 cells, 2h BIO 22h washout: N=29 cells; data representative of 3 biological repeats (n=3). Cells analysed for DMSO control, 2h BIO and 24h BIO treatment conditions are also included in (**Figure 6O, Q-R**) quantifications.

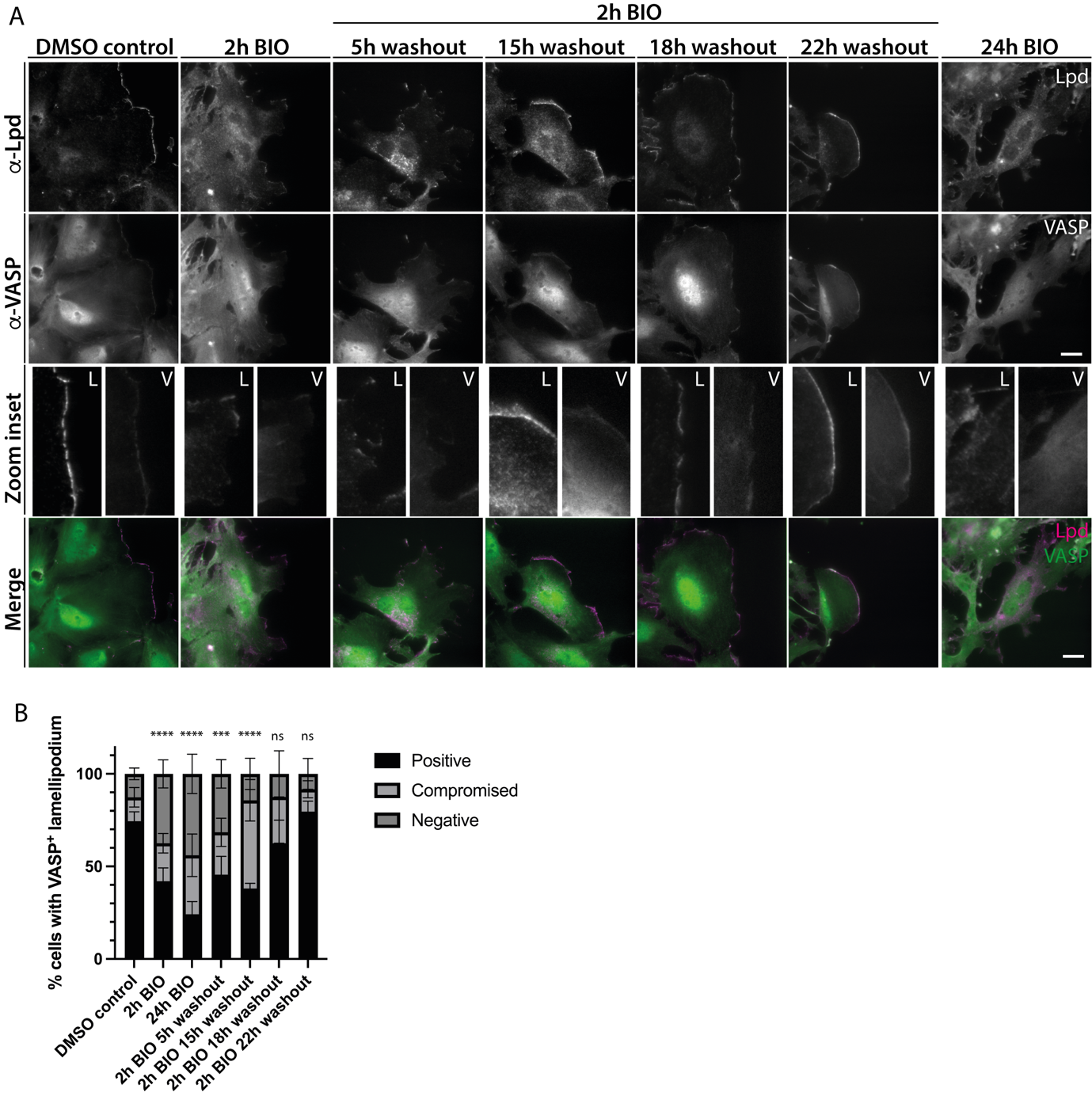

**Supplementary Figure 7. GSK3 is required for the rapid and reversible regulation of Lpd-VASP localisation in primary neural crest cells. (A)** Lpd and VASP co-immunostaining of fixed migratory cranial neural crest cells, cultured from E8.5 WT embryos, treated with DMSO control or GSK3 inhibitor, BIO, for 2 hours. Following 2h BIO treatment, explants were either fixed, or subjected to a drug wash-out whereby fresh media was applied to the explants which were allowed to recover for 5h, 15h, 18h or 22h prior to fixation. A 24h BIO treatment was also included. **(B)** Quantification of the percentage neural crest cells with a VASP-positive lamellipodium, plotted as mean ± SEM, **** p < 0.0001, ns: non-significant, chi-squared test. DMSO control: N=146 cells, 2h BIO: N=104 cells, 24h BIO: N=96 cells, 2h BIO 5h washout: N=54 cells, 2h BIO 15h washout: N=39 cells, 2h BIO 18h washout: N=36 cells, 2h BIO 22h washout: N=63 cells; data representative of 3 biological repeats (n=3).
