## Supplementary Movie Legends for "GSK3 and Lamellipodin balance lamellipodial protrusions and focal adhesion maturation in mouse neural crest migration"

**Supplementary Movie 1. Whole-mount live imaging of *Wnt1::cre* lineage-labelled delaminating neural crest cells.** Movie links to the stills shown in (**Figure 1E**). Live confocal imaging of the neural plate border (left) and cranial neural crest cells migrating towards pharyngeal arch 1 (PA1), cropped from adjacent to the neuroepithelium. Cells were imaged from a 6 somite stage, *Wnt1::cre*; *Rosa26R*^MTMG^ embryo mounted laterally. A broad cell projection (lamellipodia) is protruding out from the neuroepithelium. Maximum intensity projection movie taken over 10 minutes (1 frame/ 20 seconds), z-depth 24.5 µm, 0.5 µm per slice. Scale bar: 10 µm.

**Supplementary Movie 2. Whole-mount live imaging of *Wnt1::cre* lineage-labelled delaminating neural crest cells.** Movie links to the stills shown in (**Figure 1F**). Live confocal imaging of the neural plate border (left) and cranial neural crest cells migrating towards PA1, cropped from adjacent to the neuroepithelium. Cells were imaged from a 6 somite stage, *Wnt1::cre*; *Rosa26R*^MTMG^ laterally-mounted embryo. A broad cell projection can be seen protruding out from the neuroepithelium, as well as a post-delamination dividing cranial neural crest cell. Maximum intensity projection movie taken over 10 minutes (1 frame/ 20 seconds), z-depth 24.5 µm, 0.5 µm per slice. Scale bar: 10 µm.

**Supplementary Movie 3. Whole-mount live imaging of *Wnt1::cre* lineage-labelled delaminating neural crest cells.** Movie links to the stills shown in (**Figure 1G**). Live confocal imaging of the neural plate border (left) and cranial neural crest cells migrating towards PA1, cropped from adjacent to the neuroepithelium. Cells were imaged from a 6 somite stage, *Wnt1::cre*; *Rosa26R*^MTMG^ laterally-mounted embryo. This movie shows dynamic filopodia emanating from the edge of broader cell protrusions. A rounded cell can also be seen dividing (top left of field). Maximum intensity projection movie was taken over 30 minutes (1 frame/ 45 seconds), z-depth 40µm, 0.5 µm per slice. Scale bar: 10 µm.

**Supplementary Movie 4. Pharmacological inhibition of GSK3 reduces the speed of cranial neural crest migration *ex vivo.*** Movie links to 18 hour time-lapse stills in (**Figure 2D-F**). 10x time-lapse imaging of cranial neural crest cells migrating away from wildtype explant cultures, treated with DMSO vehicle control or the GSK3 inhibitors, BIO or CHIR99021, for 1 hour prior to imaging. Movies are overlain with dot-and-line tracks of 10 representative cells over the course of time-lapse imaging. 18 hour movie (1 frame/ 5 min). Scale bar: 100 µm.

**Supplementary Movie 5. Pharmacological inhibition of GSK3 prevents lamellipodia formation in cranial neural crest cells *ex vivo.*** Movie links to pseudo-coloured time projection stills in (**Figure 2M-O**). 60x time-lapse imaging of cranial neural crest cell protrusion dynamics, cultured from E8.5 *Wnt1::cre*; *LifeAct-EGFP* embryos. The explants were treated with DMSO vehicle control or the GSK3 inhibitors, BIO or CHIR99021, 2 hours prior to imaging, which was then completed 2-4 hours following treatment. To visualise the membrane protrusions, the actin filaments of neural crest cells were genetically labelled with LifeAct-EGFP. 10 min movie (1 frame/ 5 sec). Scale bar: 20 µm.

**Supplementary Movie 6. Genetic deletion of *Lpd* reduces the speed and persistence of cranial neural crest cells *ex vivo.*** Movie links to 18 hour time-lapse stills in (**Figure 3A-C**). 10x time-lapse imaging of cranial neural crest cells migrating away from *Wnt1::cre*; *Lpd*^+/+^ (WT), *Wnt1::cre*; *Lpd*^+/fl^ (HET) and *Wnt1::cre*; *Lpd*^fl/fl^ (HOM) explant cultures. Movies are overlain with dot-and-line tracks of 10 representative cells over the course of time-lapse imaging. 18 hour movie (1 frame/ 5 min). Scale bar: 100 µm.

**Supplementary Movie 7. Genetic deletion of *Lpd* prevents lamellipodia formation in cranial neural crest cells *ex vivo.*** Movies link to pseudo-coloured time projection stills in (**Figure 3H-J**). 60x time-lapse imaging of cranial neural crest cell protrusion dynamics, cultured from *Wnt1::cre*; *Lpd*^+/+^; *LifeAct-EGFP* (WT), *Wnt1::cre*; *Lpd*^+/fl^; *LifeAct-EGFP* (HET) and *Wnt1::cre*; *Lpd*^fl/fl^; *LifeAct-EGFP* (HOM) E8.5 embryos. To visualise the membrane protrusions, the actin filaments of neural crest cells were genetically labelled with LifeAct-EGFP. Note membrane ruffling in *Lpd* heterozygous neural crest cells, and the dynamic filopodia on *Lpd* homozygous knockout neural crest cells. 10 min movies (1 frame/ 5 sec). Scale bar: 20 µm.
